# From surfacing to stranding: The origins and dispersal dynamics of a neustonic siphonophore

**DOI:** 10.1101/2025.08.13.669909

**Authors:** River B. Abedon, Mary Beth Decker, Casey W. Dunn, Samuel H. Church

**Affiliations:** Yale University, Department of Ecology and Evolutionary Biology, New Haven, Connecticut, USA; New York University, Department of Biology, New York, New York, USA

## Abstract

The siphonophore *Physalia physalis* regularly strands along the US East Coast, yet the dynamics driving its seasonal and geographic distribution in this region remain poorly understood. Building on a new understanding of *Physalia* population structure from genomic analyses, we integrate iNaturalist observational data with a biologically-informed particle-tracking model to investigate the origins, dispersal, and stranding dynamics of *P. physalis* in the northwest Atlantic. We identify adult and juvenile observations of stranded *P. physalis* and initialize particles based on juvenile distributions. Our particle-tracking model incorporates recent research on *Physalia* drift hydrodynamics and high-resolution environmental data to simulate dispersal over twelve months. Results show a winter juvenile peak within a constrained nursery region encompassing the Straits of Florida and Gulf of Mexico. Simulated *P. physalis* undergo rapid Gulf Stream transport, with limited nursery region recirculation, indicating a source-sink population dynamic. Large-scale distribution of *P. physalis* is primarily governed by current-driven advection, while wind and wave processes drive nearshore transport and stranding. Using this framework, we characterize the role of the Gulf Stream and seasonal wind patterns in shaping *P. physalis* strandings in distinct US East Coast regions. We find that mesoscale eddies influence *P. physalis* transport in offshore regions and theorize that seasonal Gulf Stream dynamics, including eddy activity and shifts in current position and intensity, modulate strandings north of Cape Hatteras. Our study highlights the Gulf Stream’s critical role in shaping neustonic distributions in the North Atlantic and the broader transport dynamics that drive dispersal in global *Physalia* populations.

## Introduction

On the East Coast of the United States, the carnivorous siphonophore *Physalia physalis* strands by the thousands (Canepa et al. 2020; Torres Conde and Rodríguez Martínez 2023). From the Florida Keys to Cape Cod, lifeguards raise purple flags to warn swimmers when this venomous species appears near beaches (Klein 2023; Hilton 2025). Like other species of *Physalia*, *P. physalis* is neustonic, utilizing an enlarged float to reside at the sea-air interface and a sail-like crest atop to catch wind (Helm 2021). As a result, a combination of environmental forces, including ocean currents, winds, and related mesoscale oceanic processes, can influence both the offshore dispersal and onshore stranding patterns of these organisms (Bourg et al. 2022; Colaço Martins et al. 2024). Despite its conspicuous morphology and ecology, as well as the implications of its occurrence for human health and fisheries, the reproductive and transport processes underlying the distribution and abundance of *P. physalis* along the East Coast of the United States are poorly understood. While previous studies have shed light on the developmental biology, population structure, and drifting dynamics of *Physalia* spp., key questions remain, including the origins of the thousands of stranded *P. physalis*along the US East Coast and the physical oceanographic drivers of offshore and near-shore transport.

Recent changes in the understanding of *Physalia* diversity have made it critical to develop a clearer picture of large-scale dispersal patterns. Dozens of *Physalia* species were described through the end of the 19th century (Pugh 2019). In the 20th century, it was recognized that the group had been over-split, and all species were synonymized as a single species, *Physalia physalis* (Totton 1960). However, recent population genomic analyses found that there are four or more species with distinct ranges across the world’s oceans and clear morphological distinctions (Church et al. 2025). Substantial genetic structure within *Physalia* species was also observed on a global scale (Church et al. 2025). These subpopulations reflect distinct oceanic regions of origin, as predicted using backtracking models of drift. The species *P. physalis* has a broad range in the Atlantic Ocean, encompassing both sides of the North Atlantic basin and the western side of the South Atlantic basin (Church et al. 2025). Based on genetic markers, *P. physalis* observed on the East Coast of the United States were linked to specimens collected on the Gulf of Mexico coast and in Bermuda, and specimens with genetic linkage to this northwest Atlantic population have occasionally been found as far east as the Azores (Church et al. 2025). This subpopulation was found to be distinct from those collected in the northeastern Atlantic, including Ireland, Spain, and the Canary Islands, and its relationship to other western Atlantic *P. physalis*, which are observed stranding from the Caribbean Sea to Uruguay, is unknown (Church et al. 2025). This study examines the distribution and strandings of specimens in the northwest Atlantic, defined here as the western side of the North Atlantic basin north of the Caribbean Sea, as well as the Gulf of Mexico.

Given the seasonal progression of strandings from south to north along the East Coast of the United States, as well as individual sightings of northwest Atlantic *P. physalis* in the Azores, we hypothesize the Gulf Stream plays a major role in the dispersal of *P. physalis* in this region. The powerful western boundary current begins in the Gulf of Mexico, follows the contour of the southeastern US continental shelf, and then enters open waters past Cape Hatteras, where it widens and meanders as it flows across the North Atlantic (Hogg 1992; Meinen and Luther 2016). Western boundary currents play an influential role in shaping dispersal and stranding patterns of *Physalia* spp. (Bourg 2024). Specifically the East Australian Current acts as both a barrier to and facilitator of *Physalia* spp. stranding, depending on current dynamics, and demonstrate the role of associated eddies in facilitating cross-frontal transport of *Physalia* spp. in these western boundary regions (Bourg 2024). However, it is unclear whether similar dynamics influence *P. physalis* transport within the North Atlantic western boundary system and stranding patterns along the US East Coast. The Gulf Stream is also known to play a broader role in shaping marine ecosystems and the distribution of marine organisms in the northwest Atlantic. The current has an especially strong effect on the population structure and biogeography of taxa transported primarily by currents and wind, including pelagic plankton and surface-dwelling neuston (Yoder et al. 1981; Hare and Cowen 1996; Taylor et al. 2010; Naisbett-Jones et al. 2017). As a result, our findings on *P. physalis* dispersal in this region have broad implications for planktonic and neustonic species in the northwest Atlantic.

*P. physalis*, like all siphonophores, are colonial, composed of coordinated, physiologically-integrated, functionally-specialized zooids (Munro et al. 2019). In *Physalia* spp., these zooids are clustered on the ventral side of an inflated float called the pneumatophore, which provides buoyancy to keep the colony at the surface (Wittenberg 1960). Wind-forcing on this float and its protruding crest provides a propulsive force that has been likened to a warship, as *P. physalis’* common name, Portuguese man o’ war, references (Pugh 2019). Previous *Physalia* spp. movement studies have demonstrated the influence of the crest’s camber, which results in an angled response to the wind that decreases as wind speed increases (Totton and Mackie 1956; Iosilevskii and Weihs 2009; Lee et al. 2021; Bourg et al. 2024). Colonies are additionally either “right-handed” or “left-handed”, as they exhibit distinct left-right asymmetry which influences their orientation to the wind and affects their point of sail and long-distance trajectories (Totton and Mackie 1956; Iosilevskii and Weihs 2009; Lee et al. 2021; Bourg et al. 2024).

Although the distribution and dispersal dynamics of *P. physalis* are impacted by its reproductive biology, the specific temporal and geographic dynamics of reproduction and early development are poorly understood. Adult *Physalia* spp. colonies possess gonodendra, a compound reproductive structure containing gamete-producing gonophores and locomotive nectophores, that detach from the colony, mature independently, and release gametes of only one sex (Munro et al. 2019; Oguchi et al. 2024). However, specific reproductive dynamics are not understood, such as the depth and timing of fertilization and early development, as mature gonodendra have rarely been observed in the ocean (Totton and Mackie 1956). Developmental studies indicate that juvenile *P. physalis* surface once their float reaches sufficient size to provide positive buoyancy (Munro et al. 2019). It has been hypothesized, but not demonstrated that northwest Atlantic *P. physalis* reproduce during the autumn and have a 12-month life cycle (Ferrer and González 2021), while a correlation has been observed between water temperature and productivity patterns and the appearance of *Physalia* spp. in certain regions, further indicating seasonal cues for reproduction (Fierro et al. 2021; Colaço Martins et al. 2024). Given the genetic similarity among colonies found on the Gulf Coast of Texas and Cape Cod, we hypothesize that a relatively constrained nursery region exists where reproduction and early development occurs in this population, rather than reproduction occurring throughout the known adult range (Church et al. 2025). This raises further questions regarding the reproductive contributions of colonies found north of Florida along the US East Coast, which we hypothesize constitute a sink-population given the strongly directional flow of the Gulf Stream in this region.

In addition to the feeding gastrozooids and reproductive gonozooids, *P. physalis* colonies contain tentacular palpons, which dangle a curtain of venom-filled tentacles into the water column to capture prey (Iosilevskii and Weihs 2009; Munro et al. 2019; Damian-Serrano et al. 2021). In combination with its tendency toward mass strandings on beaches around the world, this has brought considerable attention to *Physalia* spp. due to the hazard posed to humans; its envenomation can cause severe pain and, in some cases, dangerous reactions in susceptible individuals, such that stranding events may shut down large stretches of beach for several days (Alam and Qasim 1991; Edwards and Hessinger 2000; Cazorla-Perfetti et al. 2012; Labadie et al. 2012; Cavalcante et al. 2020). Given the economic and public health risks of stranding events, hydrodynamic studies of *Physalia* spp. drift have been used to build model-based tools for predicting their dispersal and tracing their origins (Prieto et al. 2015; Headlam et al. 2020; Ferrer and González 2021; Lee et al. 2021; Macías et al. 2021; Bourg et al. 2022, 2024; Torres-Conde 2022). In addition, the regular appearances of stranded *Physalia* spp. on global beaches has resulted in a wealth of participatory-science data through the platform iNaturalist. This online social network, through which scientists and non-scientists can record observations of organisms seen in the wild, contains frequent observations of adult and occasional observations of juvenile *P. physalis* colonies (iNaturalist; iNaturalist Community 2024).

Together, these factors provide a strong foundation for developing a biologically realistic large-scale particle-tracking model of *P. physalis* in the northwest Atlantic, using previous experimental and model-based studies to inform the advection parameters of the model and observational data from iNaturalist to initialize simulated particles and validate model accuracy. To gain a complete understanding of northwest Atlantic *P. physalis* distribution from surfacing to stranding, our study extends beyond simulated trajectories to identify the geography and timing of early development in this population and assess the roles of regional wind patterns, the Gulf Stream, and its mesoscale eddy dynamics on dispersal. To achieve this, we conduct a large-scale analysis of northwest Atlantic *P. physalis* iNaturalist observations to characterize the temporal and spatial dynamics of East Coast strandings and early life stages. These data inform the initialization of simulated *P. physalis* in a Lagrangian particle-tracking model, parameterized using recent research of the hydrodynamics of *Physalia* spp. transport and incorporating high-resolution oceanic and atmospheric data from the one-year simulation period. We validate model accuracy using iNaturalist observational data and use a combination of experimental techniques, and quantitative and qualitative analyses of simulation outputs to examine the roles of ocean currents, wind, and mesoscale eddies in driving open-ocean distribution and stranding patterns on the East Coast of the United States. These results give insight into the distribution of northwest Atlantic *P. physalis* and the transport dynamics of *Physalia* spp. and have broad implications for the role of the Gulf Stream in shaping neustonic and planktonic population distributions in this region.

## Methods

### Overview

We first conducted a large-scale survey of *P. physalis* observations on iNaturalist to characterize the distribution and timing of US East Coast strandings and juvenile appearances. Particles were then initialized in a Lagrangian particle-tracking model based on the spatial and temporal distribution of juveniles identified in the iNaturalist survey. Particles were advected according to hydrodynamic functions informed by standard principles of Lagrangian ocean modeling and recent studies of *Physalia* spp. drift. The resulting model outputs were analyzed using a combination of qualitative and quantitative approaches, described below.

### Study area and time period

To investigate the large-scale dynamics of *P. physalis*, we first defined oceanographic and coastal regions of interest on which to focus our analyses.

We defined Area A as the waters and coastlines of the Caribbean Sea, Gulf of Mexico, and the East Coast of North America (Fig. 1), based on the previously-identified western Atlantic Ocean population range of *P. physalis* (Church et al. 2025). The juvenile study encompasses all iNaturalist *P. physalis* observations recorded from 7 January 2000 through 24 August 2024 within this region (iNaturalist Community 2024).

**Figure 1:**
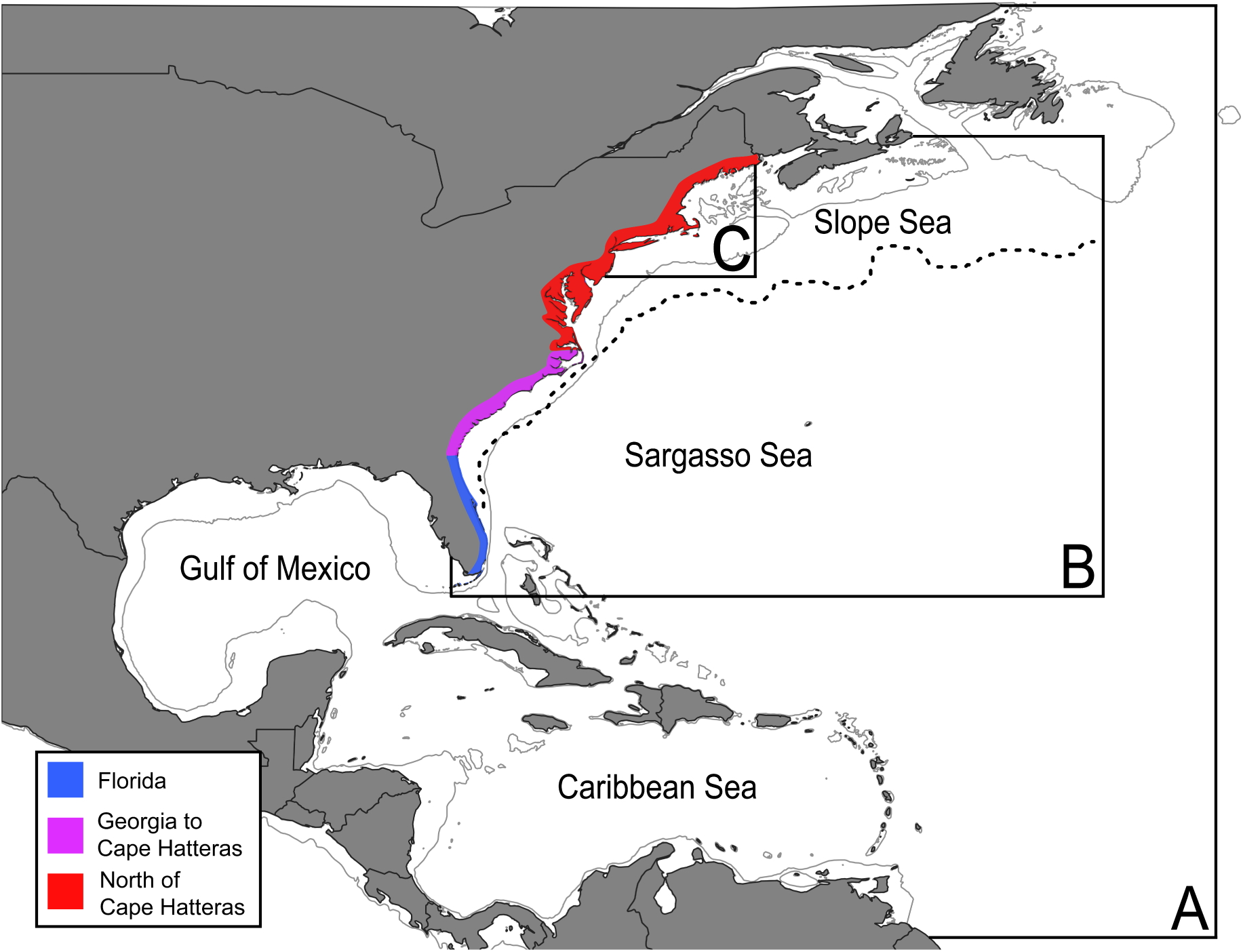
Map of study areas. Approximate boundaries of three study areas used: (A) Northwest Atlantic region within which iNaturalist observations were searched for juvenile *P. physalis* specimens. (B) Offshore region of the East Coast of the United States used in the particle-tracking study. (C) Slope and shelf region north of the Gulf Stream. Three regions of the US East Coast were also defined for analysis: Florida (blue), Georgia to Cape Hatteras (pink), North of Cape Hatteras (red). The dotted line represents the north wall of the Gulf Stream during July of 2023. The gray line indicates the 200-meter isobath.

To simulate *P. physalis* dispersal in the northwest Atlantic Ocean, we set the model domain as defined in Area A (Fig. 1). However, we focused our model output analyses on Area B, the East Coast of the United States and adjacent waters (Fig. 1), as stranding dynamics in these regions were the primary focus of the study. Likewise, to investigate observed stranding dynamics on the East Coast, we analyzed *P. physalis* observations from iNaturalist within Area B during the period from 15 January 2001 through 6 October 2024 (Fig. 1). For analysis of both simulated and observed data, we subdivided the US coastline within Area B into three coastal regions – Florida, Georgia to Cape Hatteras, and North of Cape Hatteras – based on their orientation and proximity to the Gulf Stream, which we hypothesized would influence distinct stranding dynamics (Fig. 1). We defined Area C to represent near United States waters north of the Gulf Stream offshore region (Fig. 1) and tracked particles in our simulations entering this area to investigate the dynamics of strandings in the North of Cape Hatteras coastal region, as described below.

### United States East Coast *P. physalis* observations

To understand the observed pattern of *P. physalis* strandings on the East Coast of the United States, we analyzed iNaturalist records of stranded *P. physalis* in Area B (Fig. 1) from 15 January 2001, the earliest iNaturalist observation of *P. physalis* in this region, through 6 October 2024, when data for this research was accessed (iNaturalist Community 2024). We filtered these data to exclude offshore observations as well as those from Bermuda or the Bahamas, leaving 3208 observations for analysis. We quantified the monthly observations of stranded *P. physalis* within each of the three coastal regions of interest (Fig. 1).

### Juvenile *P. physalis* observations

To characterize the distribution and timing of *P. physalis* juvenile appearances, we classified 6,818 research-grade images of *P. physalis* recorded on iNaturalist from 7 January 2000, the earliest iNaturalist observation of *P. physalis* in this region, through 24 August 2024, when data for this research was accessed, as either juvenile or non-juvenile (iNaturalist Community 2024). We determined diagnostic traits of juveniles based on *P. physalis* developmental characters (Munro et al. 2019). We classified specimens as juveniles if they had few zooids, a small float, no tentacles or a single small tentacle, and no crest or a significantly reduced crest (see supporting information, Fig. S1). We classified images in which the specimen appeared significantly damaged or in which the view of the animal was obscured as uncertain and removed them from the dataset.

To account for observer effort in the dataset, we used iNaturalist observations of three taxa of beach animals within Area A as proxies. We selected observations of hermit crabs (Paguroidea), sea stars (Asteroidea), and sand dollars and sea urchins (Echinoidea) due to the ubiquity of these animals in the area of interest and the taxonomic diversity represented (see supporting information, Fig. S2). These traits were important to ensure that the observational patterns reflected iNaturalist user effort rather than seasonal or biogeographic factors inherent to these animals. From this dataset, we created a spatial-temporal kernel density estimation (KDE) of iNaturalist user effort using the ‘gaussian_kde’ function from the SciPy statistics package. We evaluated this density function at each point in the juvenile dataset and assigned a normalization weight based on the inverse of this value (see supporting information, Fig. S2).

### Modeling juvenile distributions

To model juvenile distributions in Area A, we created a weighted spatial-temporal KDE of juvenile observations. We weighted the juvenile distribution KDE according to the normalization weight derived from the iNaturalist user effort density estimation, ensuring that observations from regions and times with lower user effort were assigned higher weights than those from regions and times with higher user effort, thereby reducing sampling bias.

We transformed the temporal variable, representing the date on which juvenile observations were made, into a day-of-year format and then subsequently converted it into a two-component circular variable using a sine and cosine function. These transformations allowed the temporal component of the juvenile distribution to be incorporated into the KDE as a continuous circular variable, preserving the cyclic nature of the observed annual patterns (see supporting information, Eq. S1 and S2).

Our starting point initialization function sampled particles from the resulting KDE, such that each generated particle starting point was assigned a location and starting date that reflected the observed juvenile distribution in space and time. The function then applied a land mask, such that particles were rejected if generated on land, and repeated the sampling process until a valid offshore location was attained. We designed the function to continue this process until 10,000 particles were generated. To ensure that particles were generated a sufficient distance from shore, we established a 50-kilometer buffer zone around land masses. If a generated particle fell within this buffer zone, the function moved the starting point to the nearest location outside the buffer zone using a breadth-first search algorithm. This method ensured that particles did not strand instantly upon initialization and are in line with the theorized open-ocean early life-stage development of *P. physalis*.

### Particle-tracking model

#### Physical data

To simulate the advection of *P. physalis* using biologically realistic parameters, we used high temporal- and spatial-resolution wind and current data from the Copernicus Marine Environmental Monitoring Service of the European Union (CMEMS) for the period 1 November 2022 through 31 October 2023. We sourced current data from the Surface and Merged Ocean Currents product of the CMEMS Global Ocean Physics Analysis and Forecast system with 1/12-degree spatial resolution and hourly temporal resolution (European Union-Copernicus Marine Service 2016). The surface *u*- and *v*-components of ocean currents in this dataset represent a merged measurement of Ekman currents at 0.494 meters beneath the surface with tidal and wave components, such that the model accounted for the effects of near-shore tidal action and Stokes drift on simulated *P. physalis* (European Union-Copernicus Marine Service 2016). We sourced wind data from the Global Ocean Hourly Sea Surface Wind and Stress from Scatterometer and Model product of CMEMS with 1/8-degree spatial resolution and hourly temporal resolution (European Union-Copernicus Marine Service 2022). These data represent the eastward- and northward-components of wind at 10 meters above the sea surface (European Union-Copernicus Marine Service 2022).

To examine the interaction between *P. physalis* and land during stranding events, we incorporated representations of landmasses within Area A during particle-tracking modeling. Polygons for all land areas in the Americas and the Caribbean at 1:50m scale were obtained from a global land database and binarized using QGIS (Natural Earth contributors; QGIS Development Team 2009).

Because the circulation model operates on a large temporal and spatial scale (Area A, Fig. 1), we accounted for the inherently stochastic nature of the ocean (Kitanidis 1994; Mohtar et al. 2018; Tewari et al. 2023). This is especially relevant in the high-energy environments represented in our model, such as the sea-air interface and the fast-moving Gulf Stream, where chaotic elements such as mesoscale eddies play a significant role in circulation dynamics (Kehl et al. 2023; Storer et al. 2023). Variables such as current, wind, ocean bottom topography, and water temperature all play a role in shaping the diffusivity constant *k* used to model this stochasticity, and it therefore varies significantly across the ocean (Groeskamp et al. 2020; Storer et al. 2023). This necessitated the implementation of a dynamic *k* based on the location of the particle. We achieved this using data from Groeskamp et al., 2020, who utilized observational data and physical modeling to calculate a full-depth global estimate of eddy diffusivity across all oceans.

To examine the direct influence of mesoscale eddies on *P. physalis* transport, we used the py-eddy-tracker Python module to detect eddies in Area B during the study period, as described below (Delepoulle et al. 2022). This module utilizes absolute dynamic topography (ADT) data to detect anomalous regions of sea surface height and define the contours of eddies according to certain criteria (Mason et al. 2014; Pegliasco et al. 2022). We used the Global Ocean Gridded L 4 Sea Surface Heights and Derived Variables product of CMEMS as the source data (European Union-Copernicus Marine Service 2021). This dataset’s 1/8-degree spatial resolution and daily temporal resolution allowed us to detect mesoscale eddies on a daily basis (European Union-Copernicus Marine Service 2021).

#### Current- and wind-driven *P. Physalis* advection

We built the *P. physalis* particle-tracking model using the Ocean Parcels toolbox (Kehl et al. 2023). The basic structure of Ocean Parcels involves the implementation of kernels to be executed on each particle in the simulation at each time step. The advection kernel defines the movement of the simulated particles in response to vector fields generated from the wind and current data. We based this function on the hydrodynamic model for *Physalia* spp. transport developed by Lee et al., 2021 and expanded by Bourg et al., 2024. This model is governed by the equations,

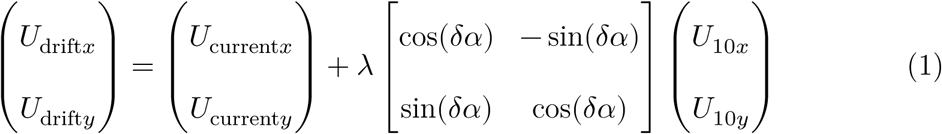

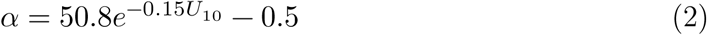

in which *P. physalis* velocity, (*U*_drift*x*_, *U*_drift*y*_), is a linear combination of the current velocity, (*U*_current*x*_, *U*_current*y*_), and a wind term. The wind response is modulated by a shape parameter, λ, which quantifies the strength of the colony’s reaction to wind forcing, or windage, and was found to be approximately 1.7% of the wind velocity measured 10 meters above the sea surface (Bourg et al. 2024). However, to account for the influence of individual variation in *P. physalis* size on a colony’s windage, our model assigned shape parameter values to particles based on a normalized distribution centered at 1.7% with a standard deviation of 0.5% (Lee et al. 2021).

*Physalia* spp. crests are cambered; therefore, the vector representing wind velocity at 10 meters above the sea surface, (*U*_10*x*_, *U*_10*y*_), is adjusted by a rotational matrix. Within the rotational matrix, the variable *α* represents the absolute value of the rotation angle as defined by Equation 2. This exponential decay function represents the observation that the intensity of *Physalia*’s angled response to wind decreases as the wind speed, *U*_10_, increases. The variable *δ* is either +1 for right-handed *Physalia* spp. or −1 for left-handed *Physalia* spp. to represent the mirrored responses to wind observed in different colony orientations (Bourg et al. 2024). During initialization, each particle is randomly assigned either a right-handed or left-handed orientation with equal likelihood, which determines the value used in the rotational matrix.

#### Stochastic advection

Our *P. physalis* particle-tracking model incorporates stochasticity by simulating random-walk using a discrete form of the Langevin equation,

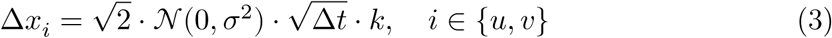

in which Δ*x*_*u*_ and Δ*x*_*v*_ are incremental changes added to a particle’s movement vector at each time step, 𝒩(0, *σ*^2^) is a Gaussian distributed random variable, Δ*t* is the time step, and *k* is the dynamic eddy diffusivity coefficient defined using data from Groeskamp et al., 2020 (Ross and Sharples 2004; Burdzy and Chen 2008; Kehl et al. 2023).

Model execution

Other than the advection kernel, we implemented three additional kernels to deactivate particles that drift outside the bounds of the model data, freeze stranded particles, and track and record strandings. Each model run, which we define as a ‘simulation’, used a time step of 6 hours, and simulation data were output every 24 hours in zarr format. The model tracked stranded particles and output their location in a comma-separated values file. We filtered all stranded particle locations to include those within Area B (Fig. 1) and in the United States to avoid the confounding effects of variable iNaturalist usage in other countries when comparing simulated data to observed data.

We ran 25 simulations for 365 days beginning on 1 November 2022, each generating 10,000 unique starting points over the course of the runtime according to the juvenile density estimation identified in Area A (Fig. 1). We chose the start date to capture the temporal peak in juvenile appearances observed in the juvenile survey and the simulation runtime to capture the seasonal pattern of *P. physalis* strandings on the East Coast observed in the iNaturalist data.

### Particle-tracking model analysis

#### Output animations

To analyze the movement of simulated *P. physalis* over time, we created model animations with geospatial visualization of nearby land masses. The animations tracked the daily positions of particles throughout the simulations. We utilized several techniques to enhance our interpretation of these animations: we overlaid a gradient-mapped representation of daily sea-surface temperature from the CMEMS Global Ocean Physics Analysis and Forecast product to visualize the Gulf Stream; we added vector representations of wind and current data, scaled to represent their influence on the simulated *P. physalis* trajectories (Eq 1); and we incorporated visualizations of mesoscale eddies detected using py-eddy-tracker, as described below (Delepoulle et al. 2022). These techniques enabled us to examine the dynamics of simulated *P. physalis* transport, as well as the effects of various environmental drivers.

#### Model validation

To validate the accuracy of the *P. physalis* particle-tracking model, we quantitatively compared the spatial and temporal distributions of the simulated stranded particles to iNaturalist observations of *P. physalis* within Area B (Fig. 1) between 1 November 2022 and 31 October 2023. We calculated the divergence between these two distributions using the earth mover’s distance (EMD), a metric that quantifies the minimal “work” required to transform one probability distribution into another (Hyun et al. 2022). By incorporating both the spatial and temporal dimensions of the data, the resulting metric favors models in which particles strand in locations and at times broadly aligned with the observed data.

To compute the EMD, we discretized the observed and simulated distributions into latitudinal and temporal bins. We divided latitude into five equal bins based on the dataset range and divided time into four equal periods over the year starting on 1 November 2022. We normalized the binned data into probability distributions summing to one. We then calculated a cost matrix to represent the spatial and temporal distances between bins and used it to calculate the EMD. We normalized this value by dividing it by the maximum possible cost, allowing for better interpretability.

We calculated the EMD for each simulated stranding distribution across 25 model runs and then calculated the average. To ensure comparability, we filtered the iNaturalist data to exclude observations not on land, as they were not representative of the stranding distribution, and those recorded far inland, which were likely artifacts of delayed uploads or observations made outside the natural setting of *P. physalis*.

#### Experimental isolation of currents from wind and waves

To assess the influence of the Gulf Stream on the large-scale transport and stranding patterns of *P. physalis*, we sought to isolate the role of ocean currents from other drivers of surface transport. To achieve this, we conducted current-only simulations in which windage and Stokes drift were removed. We define windage as the component of advection resulting from wind forcing on the above-surface crest and float of the *P. physalis* colony (Eq. 1, second term), which we removed by setting the shape parameter, λ, to 0%. We also excluded the Stokes drift component of the current field, which arises from wave orbital motion rather than geostrophic or Ekman flows (Andrews and Mcintyre 1978; Clarke and Van Gorder 2018). This left the baroclinic (geostrophic) and wind-driven (Ekman) components of the current field, which are the primary constituents of Gulf Stream flow (Stommel 1948). As with the baseline simulations, we ran 25 current-only simulations for 365 days beginning on 1 November 2022, with particles generated as described above. To ensure comparability among simulations, we set a random seed based on the run number such that starting point initialization would be identical for corresponding simulations under either condition.

We determined the number of stranding events in each baseline and current-only simulation using output data that recorded the location and timing of particles classified as stranded by the land check kernel. We averaged these values across each condition and compared them quantitatively using a Welch’s *t*-test. Additionally, we plotted stranding distributions from representative simulations under both conditions and compared them visually to identify specific regions where the removal of windage and Stokes drift altered stranding patterns.

Beyond stranding dynamics, we also sought to better understand environmental drivers of the long-term trajectories of *P. physalis*. To investigate this, we analyzed the divergence between particles with identical initializations in baseline and current-only simulations over time. We first filtered simulation outputs to identify particles that did not strand under either condition. At 15-day intervals, we calculated the distance between each particle’s location in the baseline simulations and its corresponding location in the current-only simulations. We determined the average distance for all of the filtered particles at each 15-day interval and then averaged this value across all 25 simulations to obtain a robust measure of particle trajectory divergence.

We expected the random-walk component of our model to produce compounding divergence in corresponding particle trajectories overtime, as small stochastic displacement in particle positioning may place particles with identical initializations under the influence of different environmental advection forces, resulting in amplified divergence as the simulation progresses. To isolate this influence of model stochasticity from the effect of windage and Stokes drift removal on particle trajectory divergence, we conducted a stochastic control analysis using five pairs of current-only simulations with identical initializations. We calculated the average distance between corresponding particles in these simulations at each 15-day interval, providing a measure of trajectory divergence due solely to the influence of model stochasticity. We used this as a stochastic control to better understand the role of currents in the long-term trajectory of simulated *P. physalis*. We estimated the divergence rates under both experimental and control conditions using a mixed linear model regression. The difference between these rates, captured in the condition-by-time interaction term, represents the contribution of windage and Stokes drift to particle trajectory divergence.

#### Determining drivers of strandings north of Cape Hatteras

To investigate the environmental drivers contributing to seasonal strandings of *P. physalis* in the North of Cape Hatteras coastal region (Fig. 1), we examined the conditions under which simulated particles exited the Gulf Stream and moved north toward the coast. We used entries into Area C (Fig. 1) to quantify northward advection events out of the Gulf Stream. We recorded the number of particles entering Area C for each simulation during consecutive two-week intervals over the full runtime.

To assess the influence of wind on the pattern of advection from the Gulf Stream into Area C, we determined the daily average wind velocity within the region using the Global Ocean Hourly Sea Surface Wind and Stress from Scatterometer and Model product of CMEMS (European Union-Copernicus Marine Service 2022). We plotted daily average wind values for the region and further aggregated the data into two-week intervals for direct comparison with the Area C particle entry data. We examined how both the average magnitude and the persistence of southerly wind patterns within those intervals influences the northward exit of simulated *P. physalis* from the Gulf Stream using a Pearson correlation analysis.

#### Tracking mesoscale eddies

To determine how mesoscale eddies influence the movement of simulated *P. physalis*, particularly in the offshore region of the Gulf Stream, we identified and visualized eddies within Area B (Fig. 1) throughout the simulation period. In order to achieve this, we used the Python module py-eddy-tracker (Delepoulle et al. 2022) along with satellite-derived absolute dynamic topography (ADT) data obtained from CMEMS (European Union-Copernicus Marine Service 2021). Following py-eddy-tracker’s standard procedure, we first applied a Bessel high-pass filter of 700 kilometers to the ADT data to isolate mesoscale features from large-scale sea-surface topography. We then detected anomalous regions of sea surface height using a steepness threshold greater than 0.002 meters per kilometer. We delineated the boundaries of the detected eddies, allowing for a maximum boundary-roundness shape error of 55%, and visualized those boundaries in areas of interest during each day of the study period (Mason et al. 2014; Delepoulle et al. 2022).

We integrated mesoscale eddy visualizations with our current-only simulation animations to assess their influence on particle trajectories. We used current-only simulations to ensure that the dynamics we observed were not the result of unseen wind advection. To enhance interpretability, we identified relevant eddy-driven events and filtered the dataset to include only particles located within the geographic bounds of the eddies during the corresponding timeframes. This approach allowed us to clearly visualize the dynamics of interest without visual interference from unrelated particles.

## Results

### United States East Coast *P. physalis* observations

To better understand the observed dynamics of *P. physalis* strandings on the East Coast of the United States, we analyzed monthly patterns of *P. physalis* iNaturalist observations in three coastal regions (Fig. 1).

There were frequent observations of *P. physalis* strandings from Florida to New England on iNaturalist (Fig. 2). 81.9% of all US East Coast observations during this period were in Florida (Fig. 2A). Observations of strandings in this region increased sharply from October to December and remained high throughout the winter months of January and February before gradually declining throughout the spring (Fig. 2A). Observations of Florida strandings were at their lowest during July, August, and September, with 8 of the 13 years surveyed lacking any recorded observations of *P. physalis* during these months (Fig. 2A).

**Figure 2:**
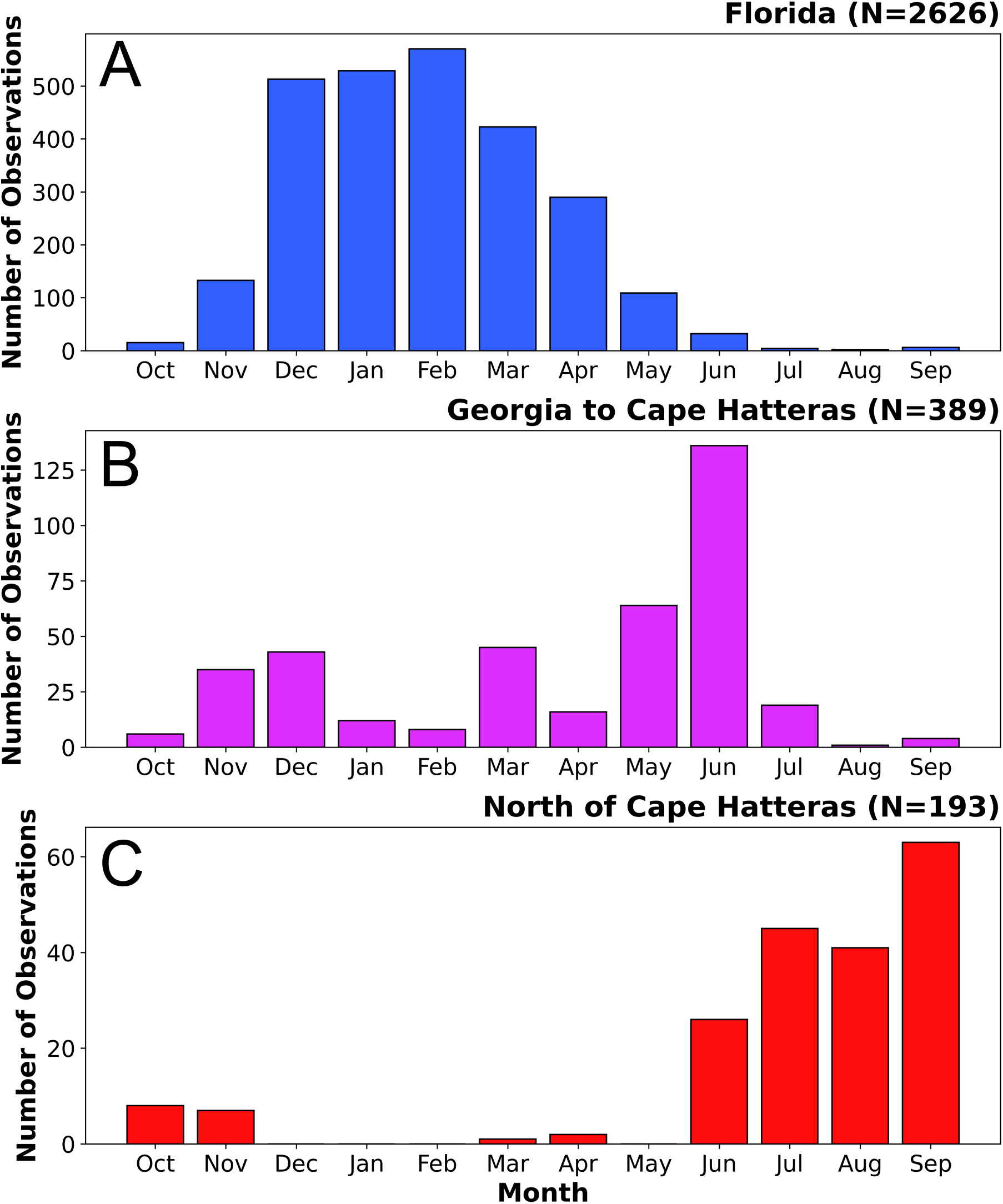
Stranding patterns on the US East Coast by region. Aggregated observations of monthly stranded *P. physalis* from iNaturalist during 2011-2024 by region of the US East Coast (Fig. 1). Note the different y-axis range in each panel.

12.1% of US East Coast *P. physalis* observations were in the Georgia to Cape Hatteras coastal region, with a pronounced peak in June (Fig. 2B). This was followed by a sharp drop in July and very few observations in August and September; as in Florida, most years surveyed lacked any *P. physalis* observations in the Georgia to Cape Hatteras region during these two months. However, the seasonal pattern in this region was less coherent and the temporal distribution in observations was more diffuse than in Florida, with smaller peaks during November, December, and March (Fig. 2B).

By contrast, the North of Cape Hatteras coastal region exhibited the strongest seasonal pattern in observations of *P. physalis* strandings (Fig. 2C). Nearly all observations in this region occurred within the six month period from June to November, with only 3 of 193 total observations recorded outside this period (Fig. 2C). Although strandings north of Cape Hatteras made up only 6% of iNaturalist *P. physalis* observations, strandings were observed at a greater rate in this region in July, August, and September compared to the other regions and were observed during these months in all but two years.

### Juvenile *P. physalis* observations

To determine the distribution and life cycle dynamics of northwest Atlantic *P. physalis* and model their origins in our particle-tracking model, we identified observations of juveniles in the iNaturalist record.

From the 6,819 observations retrieved from iNaturalist, we identified 194 as juveniles and classified an additional 86 observations as uncertain (Fig. 3A; Fig. 3B). Of the juveniles identified, we found a substantial majority, 72.1% (140 observations), are on the east coast of Florida and the Florida Keys (Fig. 3A; Fig. 3B). We found 24.2% (47 observations) of juveniles along the coast of the Gulf of Mexico, with notable concentrations along the Florida Panhandle-Southern Alabama coastline and the Texas coast (Fig. 3A; Fig. 3B). The remainder of identified juveniles were sparsely distributed further north along the US East Coast (2.6%, 5 observations) and within the Caribbean (1%, 2 observations) (Fig. 3A; Fig. 3B). We identified notable geographic outliers in these two regions, with one juvenile observed as far north as the US state of Delaware and another far to the east on the Caribbean island of Saint-Martin (Fig. 3A).

**Figure 3:**
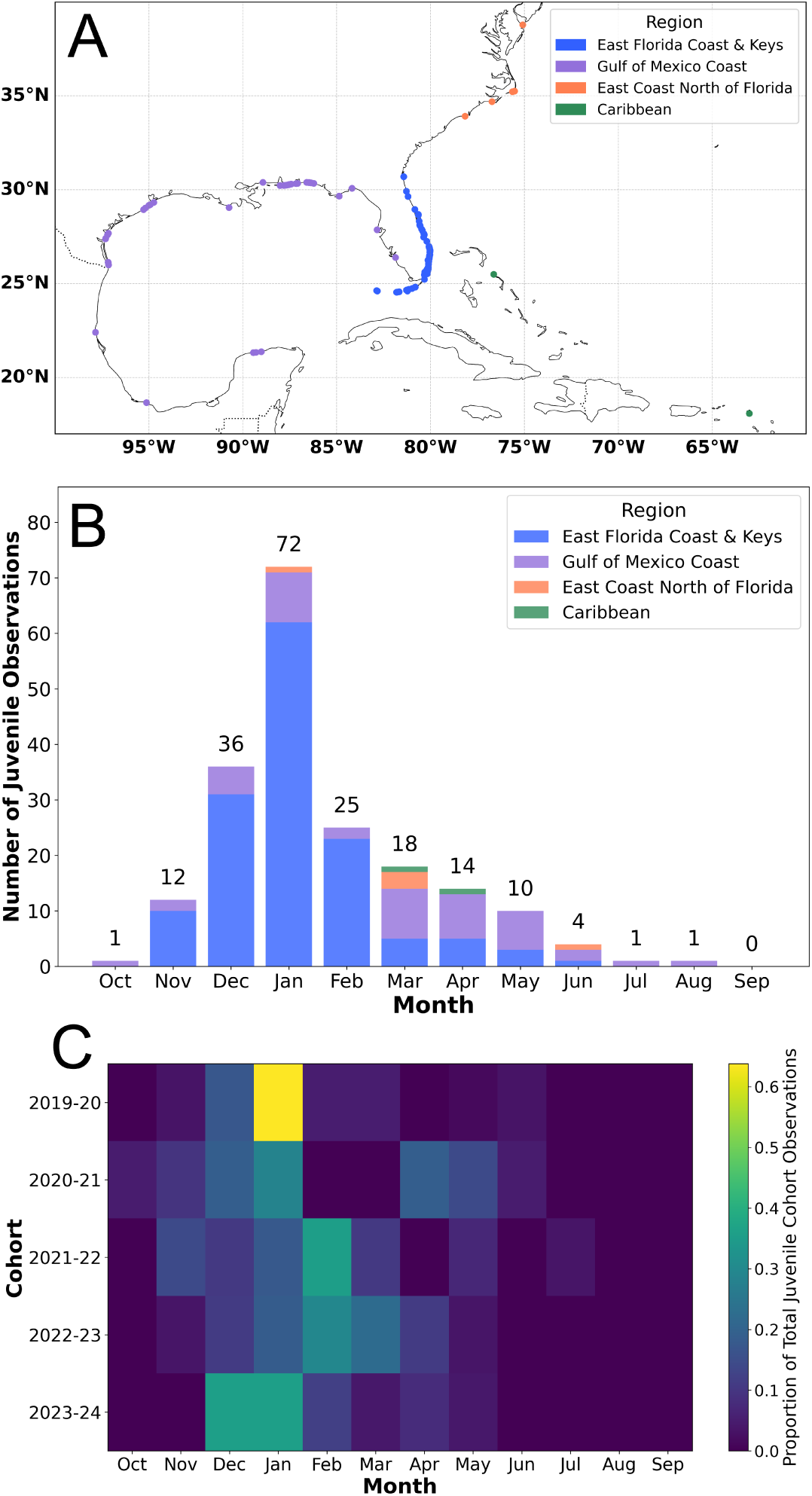
Spatial and temporal occurrences of stranded juvenile *P. physalis* observations. (A) Locations of 194 iNaturalist observations of stranded *P. physalis* identified as juveniles within Area A (Fig. 1). For simplicity, we combined observations from the Georgia to Cape Hatteras and North of Cape Hatteras regions (Fig. 1) into one East Coast North of Florida region, labeled in orange. (B) 2014 to 2024 aggregated monthly observations of stranded *P. physalis* identified as juveniles by region. (C) Proportion of total juvenile cohort observations seen in each month for all cohorts containing at least 25 identified juvenile observations (2019 - 2024).

In addition to the observed geographic patterns, we identified a distinct temporal pattern in observations of juvenile *P. physalis* (Fig. 3B). Data aggregated from the first juvenile sighting in 2014 through 2024 showed that juvenile observations peaked in December and January, accounting for 55% of total observations (Fig. 3B). We saw progressively fewer observations from February through July, with a single juvenile reported in July and August in Area A (Fig. 1), both of which were found on the Gulf of Mexico coast (Fig. 3B).

Juvenile observations on the east coast of Florida and the Florida Keys constituted a significant portion of the total observations in November, December, January, and February (Fig. 3B). Starting in March, however, this region constituted a smaller share of total juvenile observations, and observations along the Gulf of Mexico coast increased proportionally (Fig. 3B). Juvenile observations on the US East Coast north of Florida and in the Caribbean also predominantly occurred later in the year, outside the primary December-January peak (Fig. 3B).

Our analysis revealed instances where multiple iNaturalist observations submitted by the same user documented a single stranding event. When accounting for this by collapsing multiple observations made on the same day in the same region into single data points, January still retained the greatest number of observed juveniles, and the east coast of Florida and Florida Keys remained the primary location for juvenile strandings, though by a smaller margin (see supporting information, Fig. S3).

Based on these aggregated observations, we observed juvenile appearances beginning in the late fall (November), increasing rapidly in the winter (December, January), and ending by the following summer (July, August). We used this pattern to compare yearly cohorts defined as all *P. physalis* juveniles appearing between October of one year and September of the next. Although the aggregated survey included observations recorded from 2006 onwards, only cohorts from 2019 through 2024 were analyzed on an individual basis, as these cohorts contained at least 25 observations each. Our analysis of these cohorts revealed the intricacies of the seasonal pattern of juvenile observations (Fig. 3C). From 2019 to 2024, we observed that January is the peak month in juvenile appearances for 3 of the 5 cohorts (Fig. 5C). However, the 2021-2022 and the 2022-2023 cohorts deviate from this trend, exhibiting their highest juvenile proportions in February (Fig. 3C). Furthermore, we observe that the patterns of juvenile appearances vary in sharpness (Fig. 3C). While some cohorts, such as 2019-2020, display distinct peaks with a high concentration of juveniles over a short period, other cohorts, such as 2020-2021, are more diffuse and contain appearances of juveniles well into the spring (Fig. 3C).

### Model initialization

We used iNaturalist observations of three classes of organisms commonly observed at the beach to create a KDE representing iNaturalist user effort across Area A, which was used to assign a normalization weight to each juvenile observation. We used the weighted juvenile observations to construct a KDE representing the distribution of juveniles in space and time. For each of the 25 simulations, we sampled a unique set of starting locations and starting times for 10,000 particles from this KDE.

We observed particles initialized throughout the Gulf of Mexico, Straits of Florida, waters off the southeastern coast of the United States, and, to a lesser extent, the northern Caribbean Sea near the Yucatán Channel (Fig. 4A). We found the highest concentration of particles initialized near Florida (Fig. 4A). A distinct band of particle density was visible at the boundary of the 50-kilometer buffer zone that we established around coastlines (Fig. 4A). Particle initialization exhibited a unimodal temporal pattern, increasing rapidly from the start of November to its peak in January, followed by a more gradual decline throughout the spring and summer (Fig. 4B). Particles initialized near Florida were more likely to be initialized during the November to February winter period, while particles on the western side of the Gulf of Mexico or further north along the US East Coast were more likely to be generated in the spring (Fig. 4A).

**Figure 4:**
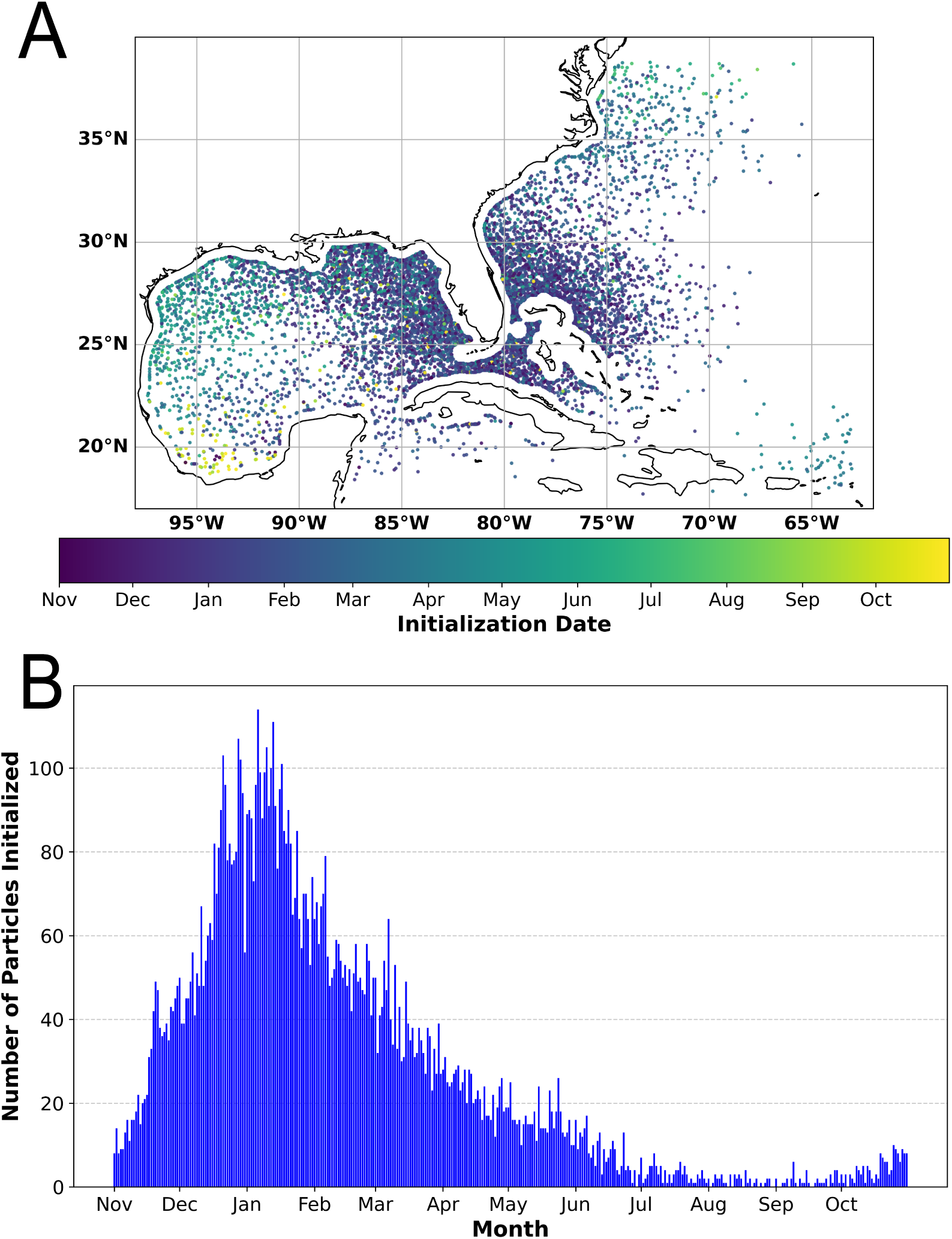
Representative location and timing of particle initialization from one simulation. (A) Distribution of 10,000 particles sampled from kernel density estimate of effort-normalized juvenile distribution. The color bar represents the start date assigned to each particle. (B) Number of particles initialized daily across the 365-day simulation period beginning 1 November 2022.

### Model validation

We simulated the dispersal of each of the 25 initialized particle distributions in the northwest Atlantic during the period 1 November 2022 to 31 October 2023 in order to model the dispersal of *P. physalis* within Area A (Fig 1). We compared the resulting stranding distributions within Area B (Fig. 1) to iNaturalist observations in the same area to validate the model.

We validated the performance of our particle-tracking model using an earth mover’s distance analysis and observed alignment between the baseline model stranding distributions and US East Coast iNaturalist observations during the same period. Across 25 simulations, each with 10,000 initialized particles run from 1 November 2022 to 31 October 2023, the average EMD between simulated strandings and observed data from the same region and period was 28.1 ± 1.8 with a maximum possible cost of 271.05, or a normalized average EMD of 0.104 ± 0.007.

In directly comparing simulated and observed *P. physalis* stranding distributions and their temporal patterns, we found key areas of alignment and divergence (Fig. 5). Overall, the spatial distributions align well, with simulated strandings occurring along similar stretches of coastline as those observed (Fig. 5A; Fig. 5B). However, in the Florida and Georgia to Cape Hatteras coastal regions, simulations generally produced a more continuous and evenly distributed pattern of strandings, whereas the observed distribution was more sporadic, with gaps and clusters of strandings (Fig. 5A; Fig. 5B). Perhaps the most notable divergence occurred in the North of Cape Hatteras region (Fig. 5). On average, only 0.9% ± 0.3% of simulated East Coast strandings occurred in the North of Cape Hatteras region, which is significantly lower than the 14.9% of total observed strandings recorded there (Welch’s *t*-test, *t*(24) = -239.9, *p* < 0.0001; Fig. 5).

**Figure 5:**
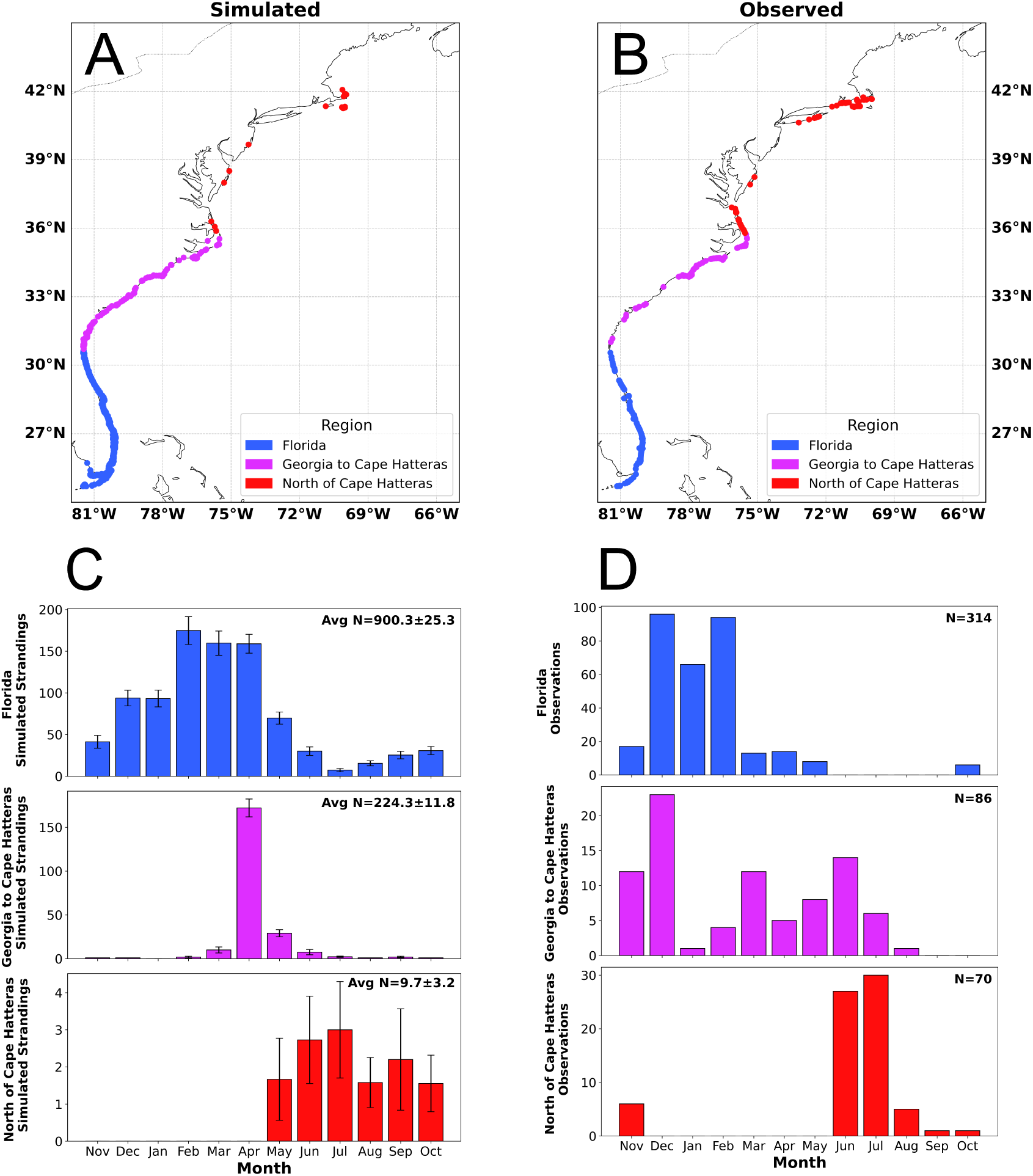
Comparison of simulated strandings with iNaturalist observations. (A) Distribution of Area B (Fig. 1) strandings from a representative simulation. (B) Distribution of iNaturalist observations of stranded *P. physalis* within Area B (Fig. 1) during the period of 1 Nov. 2022 – 1 Nov. 2023. (C) Monthly simulated strandings of particles in three coastal regions of Area B (Fig. 1) averaged across 25 simulations, including total stranding averages for each region. (D) Monthly iNaturalist observations of stranded *P. physalis* within three coastal regions of Area B (Fig. 1) during the period of 1 Nov. 2022 – 1 Nov. 2023, including total number of observations for each region.

In the Florida and North of Cape Hatteras coastal regions, we found broad alignment in the temporal stranding patterns of the simulated and observed data (Fig. 5C; Fig. 5D). In the Florida coastal region, we observed a winter peak in strandings followed by minimal activity during the summer in both the simulated and observed data (Fig. 5C; Fig. 5D). In the North of Cape Hatteras coastal region, strandings were confined to a similar time frame in both the simulated and observed data, with simulated strandings occurring from May to October and observed strandings seen from June to November (Fig. 5C; Fig. 5D). However, we observed notable divergence in the temporal pattern of strandings in the Georgia to Cape Hatteras region (Fig. 5C; Fig. 5D). While observed strandings in this region peaked in December and occurred with relative frequency through August, simulated strandings instead show a distinct April peak and comparatively fewer strandings during the rest of the year (Fig. 5C; Fig. 5D).

Despite these divergences, our earth mover’s distance analysis, along with the broad agreement between observed and simulated geographic and temporal stranding distributions, indicate that our model accurately captures the drift dynamics of *P. physalis*.

### Transport dynamics of simulated northwest Atlantic *P. physalis*

We used trajectory mapping and simulation animations to analyze the spatial patterns of simulated *P. physalis* dispersal and stranding in our particle-tracking model. To examine the environmental drivers influencing the simulated dispersal and stranding dynamics, we ran an experiment altering model parameters related to wind and waves, tracked particle exits from the offshore region of the Gulf Stream, and detected mesoscale eddies in simulation animations.

### Gulf Stream-driven transport of *P. physalis*

We used trajectory mapping to analyze the spatial patterns of simulated *P. physalis* dispersal and strandings in our particle-tracking model. Trajectories were plotted for particles in Area B that never stranded during the entirety of the model run, and for those that stranded in each of the defined coastal regions (Fig. 1).

We observed a strong influence of the Gulf Stream on the transport of simulated *P. physalis* in our particle-tracking model (Fig. 6). Across simulations, particles initialized according to our juvenile density estimation were transported rapidly northward along the US East Coast via the Gulf Stream (Fig. 6). Many particles were initialized within the direct path of the current (Fig. 6; Fig. 4). However, we also observed particles originating in the Gulf of Mexico, the Sargasso Sea, and the Caribbean Sea dispersing into the Gulf Stream (Fig. 6). This movement of particles from adjacent waters into the current system occurred consistently throughout the year-long simulation period (Fig. 6).

**Figure 6:**
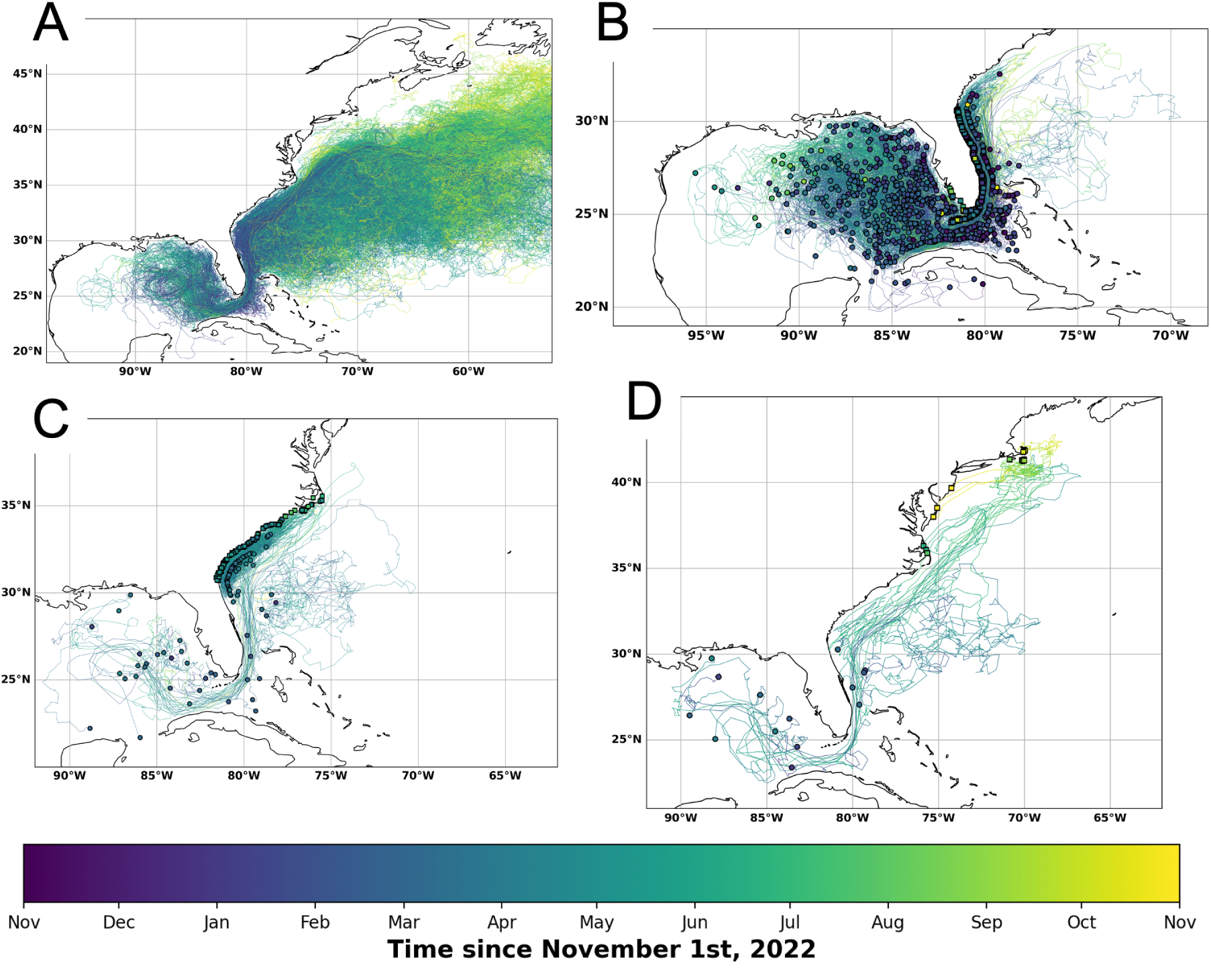
Long-distance trajectories of simulated *P. physalis* on the US East Coast. (A) Trajectories of 500 randomly selected particles from a representative simulation that entered Area B (Fig 1) and did not strand over model run. (B-D) Trajectories of all particles from the same simulation that stranded in the (B) Florida coastal region, (C) Georgia to Cape Hatteras coastal region, and (D) North of Cape Hatteras coastal region (Fig. 1). Circular points represent initialization locations, and square points represent stranding locations. The eastern limit of the map represents the eastern limit of our model domain.

Many particles that entered the Gulf Stream in our model did not strand at any point during the simulation run (Fig. 6A). Most of these particles were transported out of the Gulf Stream at some point, especially as they approached the offshore region of the current north of Cape Hatteras (Fig. 6A). Throughout most of the year, we observed that a majority of non-stranding particles dispersed into the Sargasso Sea to the southeast of the current (Fig. 1; Fig. 6A). However, between June and September, non-stranding particles in the offshore region dispersed into the Slope Sea more frequently than earlier in the year (Fig. 1; Fig. 6A). After exiting the current system, we observed particles following meandering paths that either recirculated them into the Gulf Stream, kept them within adjacent slower-moving waters through the remainder of the simulation, or eventually carried them beyond the eastern simulation boundary, where they were deactivated (Fig. 6A). A significant number of non-stranding particles, however, remained within the path of the Gulf Stream after entering this offshore region, all of which were rapidly transported beyond the easternmost simulation boundary and deactivated (Fig. 6A). Within the trajectories of particles that did not strand, we found no evidence of particles recirculating to the Gulf of Mexico after they had entered the Gulf Stream and a very limited number of particles recirculating to the Straits of Florida after they had entered the Gulf Stream (Fig. 6A).

Across 25 simulations initialized with 10,000 particles each, we observed an average of 1133.9 ± 5.3 simulated *P. physalis* stranded on the US East Coast in Area B (Fig. 1; Fig. 5A). Of the coastal regions we defined (Fig. 1), strandings were most frequent in Florida, with an average of 900.3 ± 5.2 strandings per simulation (Fig. 5C; Fig. 6B). Many of the particles stranding in Florida were initialized near the Florida coastline and stranded soon after initialization (Fig. 6B). However, we also found that a considerable number of particles that stranded in this region originated in the Gulf of Mexico and, less frequently, the Caribbean Sea; these particles dispersed into the southern Gulf Stream region, which carried them nearshore to Florida, where they subsequently stranded (Fig. 6B). While most particles traveled northward via the Gulf Stream to their stranding location on the Florida Coast, a few first dispersed into the Sargasso Sea to the east of the current and then were recirculated back to the southern Gulf Stream region before stranding (Fig. 6B). Additionally, we observed a small group of particles initialized north of Florida in the slower-moving shelf waters between the Gulf Stream and the coast, known as the South Atlantic Bight, which then dispersed south to strand along Florida’s northeast coast (Fig. 6B).

Florida strandings were frequent whenever particles were initialized nearshore or entered the southern Gulf Stream region (Fig. 6B). We observed a decrease in Florida strandings in spring and summer, coinciding with lower particle densities in the Gulf Stream and Gulf of Mexico, followed by a subsequent increase in the fall as particle densities rose again (Fig. 5C; Fig. 6B).

An average of 224.1±2.4 simulated strandings occurred in the Georgia to Cape Hatteras coastal region (Fig. 5C). As in Florida, stranded particles in this region originated from both nearby and distant source locations (Fig. 6C). Across simulations, most of the particles that stranded in this region were initialized near its coastline and accumulated in the South Atlantic Bight to the west of the Gulf Stream before ultimately being brought onshore during a mass stranding event in April (Fig. 5C; Fig. 6C). A smaller subset of particles that stranded in this region originated in the Sargasso Sea, Gulf of Mexico, and Straits of Florida, and were brought to the Georgia to Cape Hatteras region via the Gulf Stream (Fig. 6C). While some of these particles, particularly those that stranded near Cape Hatteras, moved directly onshore from within the Gulf Stream, the majority first dispersed into the South Atlantic Bight, where they sometimes remained for several months before stranding (Fig. 6C).

The North of Cape Hatteras region had the fewest simulated *P. physalis* strandings of the three coastal regions we defined, with an average of 9.6 ± 0.6 strandings (Fig. 5C). Although infrequent, simulated strandings in this region exhibited strong seasonality, occurring consistently across simulations between May and October (Fig. 5C; Fig. 6D). Particles stranding in this region had southerly origins in the Gulf of Mexico, Sargasso Sea, South Atlantic Bight, or within the path of the Gulf Stream (Fig. 6D). Despite the initialization of hundreds of particles north of Cape Hatteras according to the juvenile density estimate (Fig. 4), we did not observe any of these particles strand in the North of Cape Hatteras coastal region (Fig. 6D).

While we observed a few isolated cases of particles moving through the Pamlico Sound to the west of Cape Hatteras and stranding on the Outer Banks of North Carolina, the vast majority of particles that stranded in this region first passed through the offshore region of the Gulf Stream (Fig. 6D). Starting in May, particles in this offshore region began dispersing northward out of the current into the Slope Sea and the adjacent shelf waters of the Mid-Atlantic Bight (Fig. 6A; Fig. 6D). We observed many of these particles remaining in this region north of the Gulf Stream for several months, with the majority never stranding (Fig. 6A; Fig. 6D). However, a subset continued to disperse north and stranded on Long Island and in southern New England, while others dispersed southwest and stranded from Cape Hatteras to New Jersey (Fig. 6D).

### Impact of wind and ocean currents on simulated *P. physalis* transport

To assess the relative contribution of surface wind- and ocean current-driven forces to the long-distance transport and stranding of *P. physalis* in the northwest Atlantic Ocean, we repeated the simulations under a current-only condition by removing windage and Stokes drift and compared the resulting stranding patterns and trajectories of simulated particles to the baseline simulations.

The removal of windage and Stokes drift had a large influence on the total number of simulated strandings. Across 25 simulations each with 10,000 initialized particles run from 1 November 2022 to 31 October 2023, the average number of simulated strandings on the US East Coast was 1133.9±5.3 (Fig. 7A). Across the same number of simulations with windage and Stokes drift removed, the average number of simulated strandings in the U.S. East Coast was 354.4 ± 3.8 (Fig. 7B). Simulations without wind had 68.7% less strandings on average than simulations with wind (Fig. 7). This difference was statistically significant (Welch’s *t*-test, *t*(43.5) = 120.1, *p* < 0.0001).

**Figure 7:**
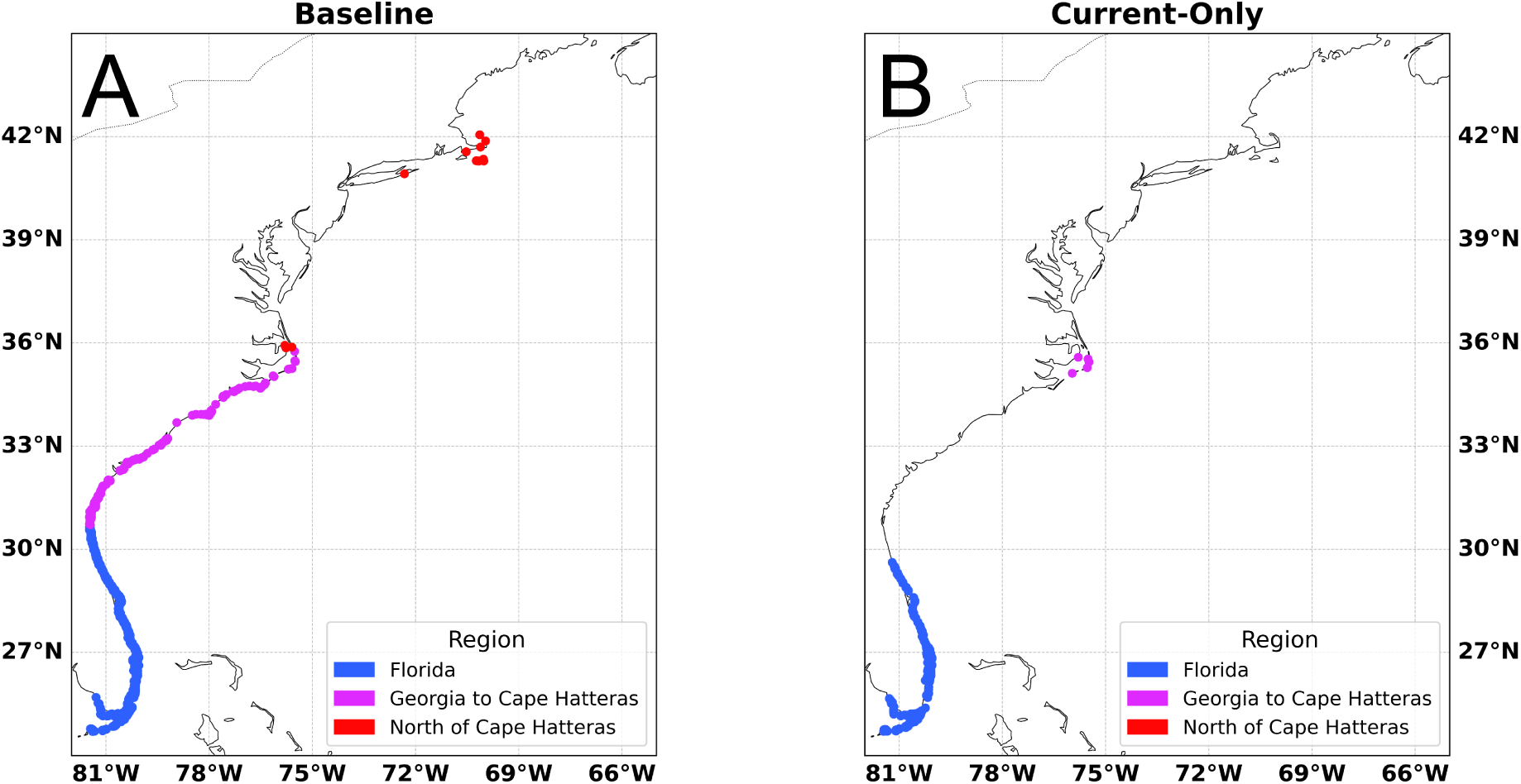
East Coast strandings of simulated *P. physalis* in (A) baseline and (B) current-only simulations. Stranding distributions of simulated *P. physalis* from representative baseline (A) and current-only (B) simulations with identical initializations. Stranding locations are colored based on previously-defined US East Coast regions (Fig. 1).

Despite this large effect, strandings persisted in certain areas under the current-only condition (Fig. 7B). In the Florida coastal region, the number of strandings remained comparatively high in current-only simulations, averaging 349.2±3.9 strandings, most of which we observed in southerly regions of the coastline (Fig. 7B). We observed a similar pattern around Cape Hatteras, where strandings occurred with relative frequency in the current-only simulations (Fig. 7B). The Georgia to Cape Hatteras region experienced an average of 5.24 ± 0.5 simulated strandings, all of which we observed on shorelines near Cape Hatteras (Fig. 7B).

By contrast, strandings in the current-only simulations were entirely absent along the coasts of northern Florida, Georgia, and the Carolinas south of the Outer Banks (Fig. 7B). We similarly observed no strandings in the North of Cape Hatteras coastal region across 25 current-only simulations (Fig. 7B).

Despite its large effect on stranding patterns, the removal of windage and Stokes drift from the model had a relatively small effect on large-scale open water transport (Fig. 8). Between the baseline and current-only simulations, average divergence of corresponding particle trajectories initially increased gradually, accelerated between days 45 and 150, and then slowed again while continuing to rise through the remainder of the simulations (Fig. 8). By the end of the simulations, particles under the current-only condition were an average of 1048.8 ± 8.3 kilometers away from their counterparts in the baseline simulations, despite having identical initializations (Fig. 8). However, we observed a very similar pattern of trajectory divergence in the control simulations, where corresponding particles diverged an average of 981.8 ± 15.5 kilometers from their counterparts, despite identical initializations and advection parameters (Fig. 8). This indicates that a large majority of divergence was due solely to stochasticity while windage and Stokes drift contributed only 67.0 ± 3.8 kilometers of additional divergence to simulated *P. physalis* trajectories over the course of a year – approximately 6.4% of the total – or 0.19 kilometers of divergence per day (mixed linear model regression, *z* = -2.30, *p* = 0.22; Fig. 8).

**Figure 8:**
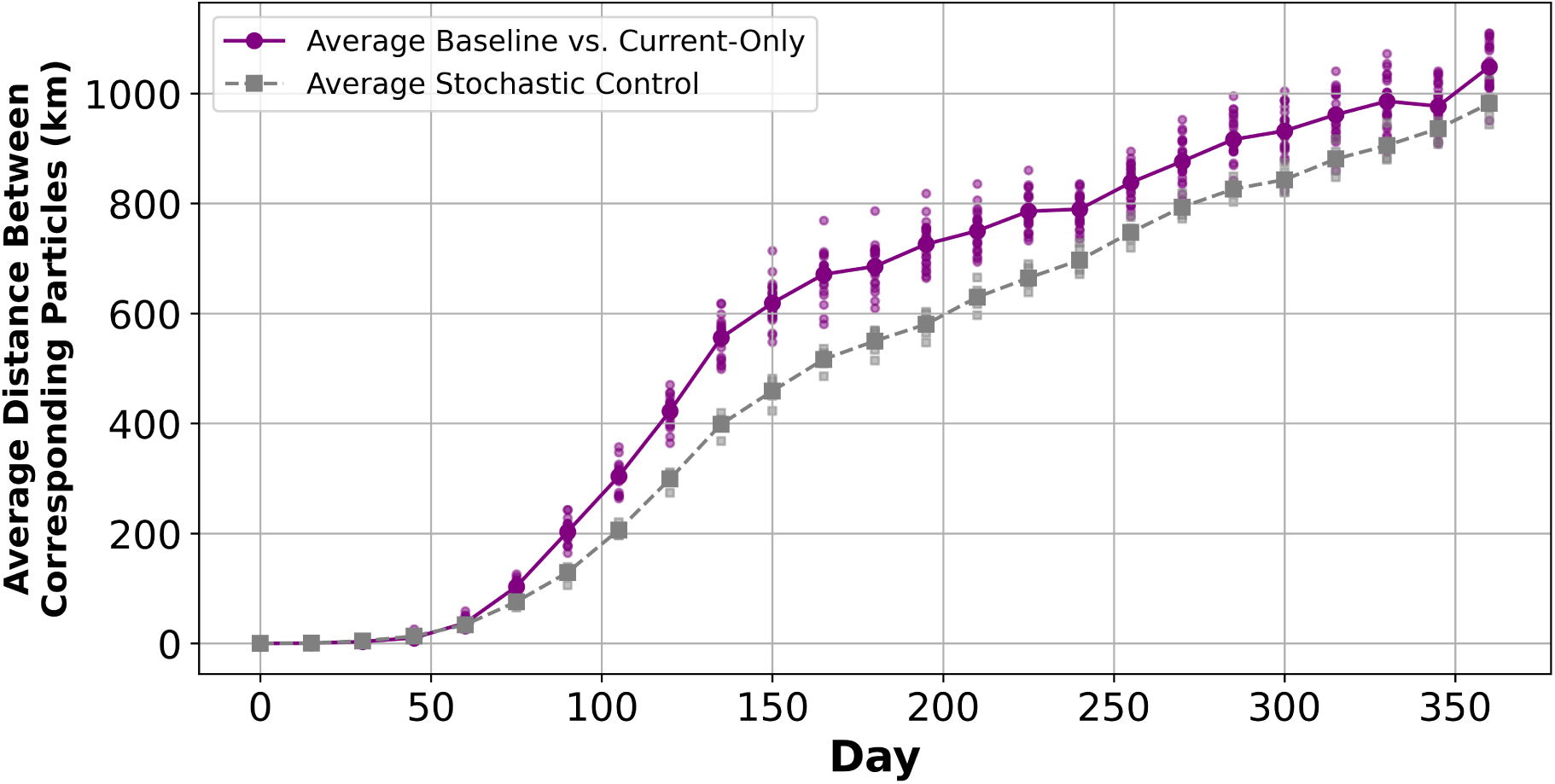
Effect of removing wind and waves on trajectories of simulated *P. physalis*. The purple dots indicate the average distance between corresponding particles with identical initializations in 25 baseline and current-only simulations is shown at 15-day intervals. The gray squares indicate the average distance between particles in ten current-only simulations, demonstrating the influence of stochasticity on corresponding particle divergence.

Animations also revealed that large-scale distribution patterns in current-only simulations were similar to the baseline. We observed that many simulated *P. physalis* approached similar coastal areas along the US East Coast in both the baseline and current-only simulations. However, far fewer particles in the current-only simulations made it onshore compared to the baseline (Fig. 8).

### Environmental drivers of simulated *P. physalis* strandings north of Cape Hatteras

Our simulations indicated that the pronounced seasonal pattern of *P. physalis* strandings in the North of Cape Hatteras coastal region (Fig. 2; Fig. 5D) coincided with the northward dispersal of particles from the offshore region of the Gulf Stream into the Slope Sea starting in late April (Fig. 6A; Fig. 6D). We further investigated this dynamic by examining the environmental drivers of particles entering the Slope Sea in our simulations.

Simulated *P. physalis* began exiting northward from the Gulf Stream in late April, indicated by a sharp increase in entries into Area C during late June and early July (Fig. 9). Strandings began shortly after initial particle entry into Area C, starting in some simulations in early May and continuing through early October (Fig. 9). Although strandings continued, the average number of particles entering Area C declined from July through October (Fig. 9).

**Figure 9:**
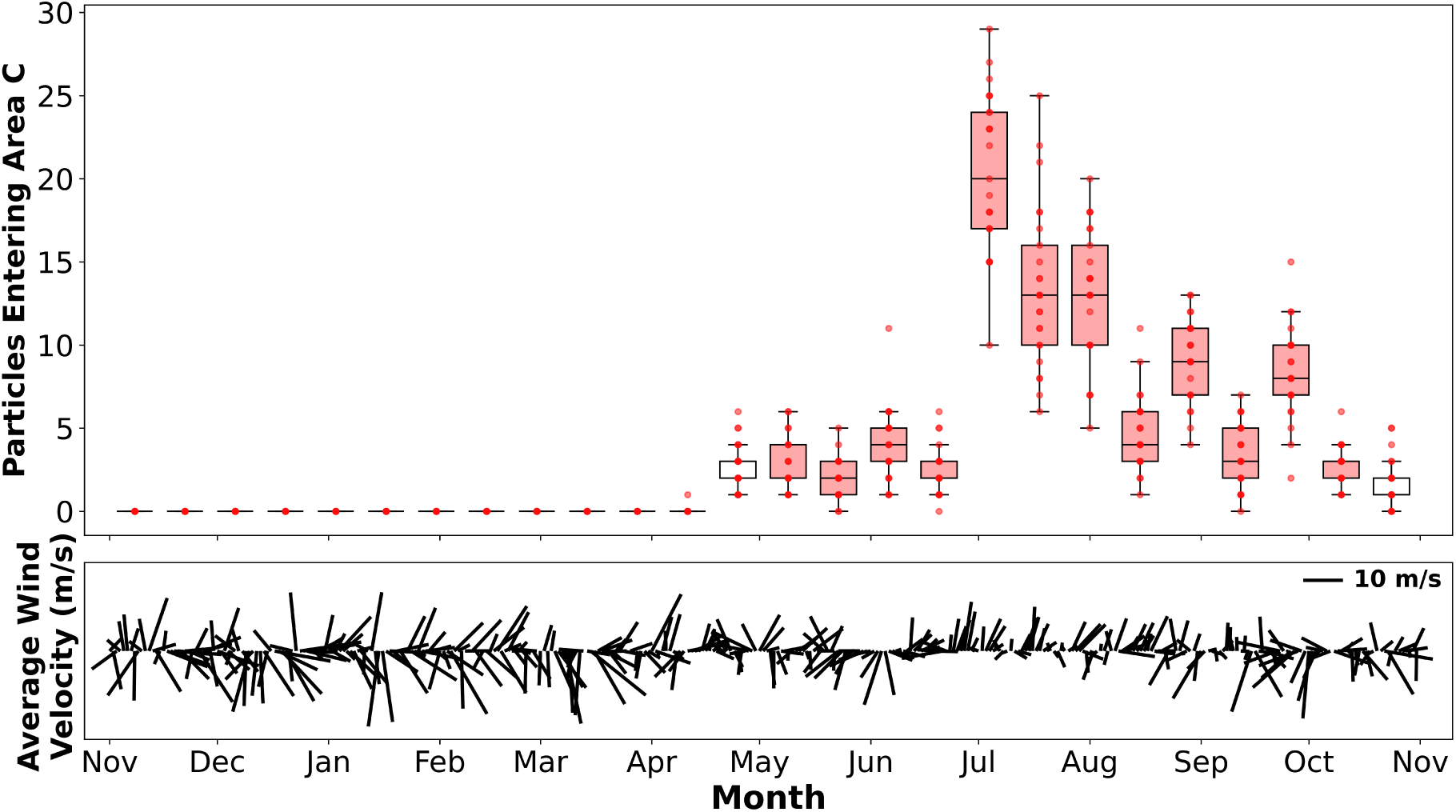
Regional wind patterns drive North of Cape Hatteras *P. physalis* strandings. Top panel shows the number of particles entering Area C (Fig. 1) over consecutive two-week intervals across model run. Red points show each simulation count, with corresponding box plots averaging the distribution across all 25 simulations. Box plots are shaded red for intervals during which at least one particle stranded in the North of Cape Hatteras coastal region (Fig. 1). Bottom panel shows the average daily wind velocity across Area C and the offshore region of the Gulf Stream in meters per second. The quiver key on the top right shows the length representing 10 m s^−1^.

While we observed intermittent southerly winds throughout the year, there was substantial short-term variability in wind direction from November to May in Area C and the offshore region of the Gulf Stream (Fig. 9). Starting in June, however, we observed a consistent southerly average wind pattern in this region, with almost uninterrupted daily southerly winds lasting for two months (Fig. 9). This period coincided with the sharp increase in the number of particles that entered Area C in late June and early July (Fig. 9).

To quantify the relationship between wind and the northward advection of particles out of the Gulf Stream, we tested the correlation between the average northward component of wind and particle entry into Area C during two-week intervals from 1 November 2022 to 31 October 2023. We found a statistically significant but moderate correlation (Pearson’s correlation, *r* = 0.412, *df* = 24, *p* < 0.05,), indicating that strong southerly winds were associated with increased entry into Area C. However, we found a stronger correlation when considering the number of consecutive southerly wind days within each two-week interval (Pearson’s correlation, *r* = 0.629, *df* = 24, *p* < 0.005,). This suggests that the persistence of southerly wind conditions, rather than the strength of southerly wind magnitudes alone, was a more robust predictor of entry into Area C. These results highlight wind as a key but incomplete environmental driver of the seasonal transport of *P. physalis* northward out of the Gulf Stream.

### Influence of mesoscale eddies on simulated *P. physalis* transport

To further investigate environmental drivers of *P. physalis* transport dynamics in the northwest Atlantic, we detected and visualized mesoscale eddies in the region using absolute dynamic topography (ADT) data and the py-eddy-tracker Python module.

Our visualization of daily mesoscale eddies in Area B (Fig. 1) throughout the simulation illustrates how the formation of eddies contributes to the movement of simulated *P. physalis* out of the Gulf Stream (Fig. 10). Many particles in our simulations dispersed out of the Gulf Stream either within the boundary of an eddy or in the peripheral currents surrounding an eddy (Fig. 10). We observed this dynamic with the warm-core eddies formed on the northside of the Gulf Stream and in the cold-core eddies formed on the southside of the Gulf Stream (Fig. 10). While this dynamic was especially prevalent in the meandering offshore region of the Gulf Stream (Fig. 10), we also observed it to a lesser extent with frontal eddies in the southern parts of the current off the Florida and Georgia to Cape Hatteras coastal regions. Cold-core eddy formation on the southern side of the Gulf Stream contributed to a significant movement of simulated *P. physalis* into the Sargasso Sea, while warm-core eddy formation on the northern side of the Gulf Stream contributed to a lesser, but still significant movement of simulated *P. physalis* into the Slope Sea (Fig. 10). A portion of the simulated *P. physalis* that dispersed into these areas recirculated back into the Gulf Stream, while others remained in the waters to the north and south of the current for the remainder of the simulation (Fig. 6A; Fig. 10).

**Figure 10:**
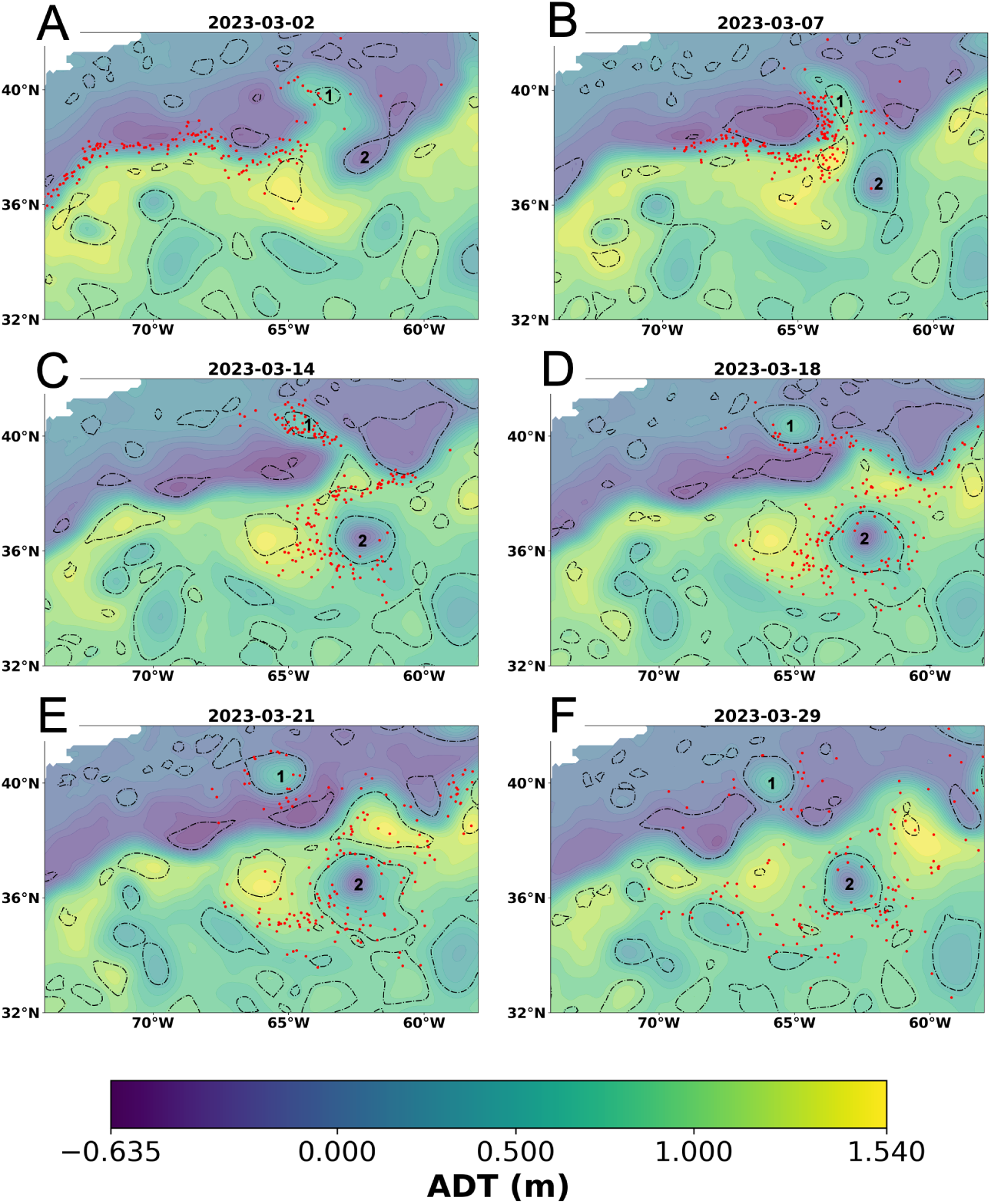
Influence of eddies on simulated *P. physalis* dispersal in the Gulf Stream offshore region. Particle transport during formation of two eddies in the Gulf Stream from representative current-only simulation. Dotted lines indicate eddy contours, color gradient indicates absolute dynamic topography (ADT) in meters, red points indicate particles, selected from a current-only simulation and filtered for particles relevant to eddy dynamics (see Methods). From early to mid-March, a pronounced Gulf Stream meander forms and pinches off into a warm-core eddy (clockwise rotation) to the north and a cold-core eddy (counterclockwise rotation) to the south (A–B). The warm-core eddy (1) moves northwestward, carrying particles into the Slope Sea, while the cold-core eddy (2) moves southward, dispersing particles into the Sargasso Sea (C–E). As both eddies expand and evolve, some particles are recaptured by the Gulf Stream, but many continue drifting west or south by eddy advective processes. By late March, most particles have exited direct eddy influence but remain displaced from the Gulf Stream (F).

## Discussion

Our findings suggest that *Physalia physalis* in the northwest Atlantic Ocean constitute a single population that experiences a seasonal winter peak in juveniles within a nursery region encompassing the Straits of Florida and Gulf of Mexico. Colonies subsequently disperse northward along the East Coast of the United States via the Gulf Stream, populating much of the current by the early spring, and do not recirculate into the nursery region, suggesting source-sink dynamics in this population. Seasonal changes in dominant wind patterns and Gulf Stream dynamics influence the movement of colonies out of the current, resulting in distinct seasonal stranding patterns in different regions of the US East Coast influenced by their orientation to and distance from the Gulf Stream. Our findings also highlight the critical role of current-driven advection in the long-distance transport of *P. physalis*, and of wind in driving near shore transport and stranding events.

The distribution we identified of juvenile *P. physalis* observations on iNaturalist shows that northwest Atlantic *P. physalis* juveniles are primarily observed on the shores of the Straits of Florida and Gulf of Mexico, with rare observations further north along the US East Coast (Fig. 3A). Our particle tracking results further support the Straits of Florida and Gulf of Mexico during the winter and early spring as the likely area and time of origin for the northwest Atlantic *P. physalis* population, showing that particles initialized in this region and period and modeled according to biologically realistic parameters produce stranding distributions on the US East Coast that closely align with observed stranding patterns in both space and time (Fig. 5; Fig. 6). The misalignment in the timing of simulated and observed strandings in the Georgia to Cape Hatteras coastal region, which we theorize resulted from a premature accumulation of particles initialized in the South Atlantic Bight region (Fig. 5C; Fig. 5D; Fig. 6C), suggests our simulations may overestimate the South Atlantic Bight as a juvenile *P. physalis* surfacing location, and further supports the Straits of Florida and Gulf of Mexico as the most likely regions of origin. Because the outliers were observed further north on the US East Coast were observed after the average peak in juvenile abundance (Fig. 3B), we hypothesize that they originated further south and were transported north by the Gulf Stream during early development (Fig. 3A). Additionally, while our simulations show a small number of particles initialized in the northern Caribbean Sea stranding along the US East Coast in alignment with the observed distribution (Fig. 6B; Fig. 6C), the absence of *P. physalis* juvenile iNaturalist observations in this region leads us to consider it a less likely region of origin for this population (Fig. 3A). These findings support our hypothesis that reproduction and early development in the northwest Atlantic population of *P. physalis* occurs within a relatively constrained nursery region and reject the alternative hypothesis that successful reproduction occurs throughout the known adult range of *P. physalis*.

Observations of both adult and juvenile *P. physalis* on the East Coast of Florida peak from December to February (Fig. 2A; Fig. 3B; Fig. 3C), indicating a seasonal population increase during this period. While we considered whether this pattern could be caused by physical drivers of strandings rather than reproductive dynamics, we found it unlikely given that dominant easterly winds, which would facilitate strandings on the East Coast of Florida, are present in Florida year-round (Peng et al. 1999). The timing of the seasonal population increase aligns with the winter peak in net primary productivity observed in the Gulf of Mexico’s surface waters (Muller-Karger et al. 2015; Yang et al. 2022), consistent with previous findings linking *P. physalis* occurrences to primary productivity in the North Atlantic (Colaço Martins et al. 2024).

Our simulations show that once *P. physalis* colonies enter the Gulf Stream, they do not appear to recirculate back into the nursery region within the twelve month period of our model. This supports our hypothesis that the colonies along the US East Coast constitute a population sink, unlikely to contribute to new generations (Fig. 6B). This finding is particularly notable given the near absence of adult *P. physalis* strandings in Florida during the late summer, just before the sharp rise in both adult and juvenile observations in the fall; over the 13-year study period, only six *P. physalis* observations were recorded in Florida in July and August (Fig. 2A; Fig. 3B), with similarly few observations during this period elsewhere along the Gulf Coast. It is possible that only a small number of adult *P. physalis* ultimately contribute to the reproductive season each year. However, we present two alternative mechanisms by which adults present in the nursery region during a reproductive season might contribute to the subsequent reproductive season despite low population numbers in the preceding months. The first is delayed fertilization, in which detached gonodendra persist in subsurface waters for several months after detaching from the colony, maturing and releasing gametes once conditions are optimal in the fall. The second is delayed development, in which *P. physalis* larvae undergo an extended subsurface early-development phase, surfacing once conditions are optimal in the fall. While these mechanisms are not mutually exclusive, we find the former more likely given recent evidence suggesting *Physalia* gonodendra mature after they are released (Oguchi et al. 2024). However, further research is needed to clarify the developmental timeline of *P. physalis* colonies and their gonodendra, the depth and timing of gamete release and fertilization, and the seasonal population dynamics of *P. physalis* in the nursery region.

The results of our models using wind, currents, Stokes drift and stochastic advection show broad alignment with observed stranding patterns as reported on iNaturalist (Fig. 5). We find that our simulated strandings align more closely with observed *P. physalis* strandings aggregated over multiple years than *P. physalis* strandings from 1 November 2022 to 31 October 2023, the period of simulations (Fig. 2; Fig. 4C). This may result partly from our particle initializations, which reflect a ten-year aggregate of juvenile stranding distributions. However, it also highlights certain limitations in our physical ocean modeling strategy. As discussed by Bourg et al., 2024, our advection model excludes submesoscale dynamics and other fine-scale processes not represented at the spatial and temporal resolution of our environmental datasets, as well as certain drivers of surface advection such as swell-induced Stokes drift (Bourg 2024). Given these simplifications and the inherently stochastic nature of ocean transport, it is unsurprising that our simulation results align more closely with a multi-year aggregate rather than exactly reflecting the period in which they were modeled.

Our findings using models that isolate the effects of currents from wind suggest that ocean currents in the northwest Atlantic are the primary drivers of the large-scale distribution and coastal proximity of *P. physalis* colonies, while windage and associated wave-driven processes are primarily responsible for the final step of bringing colonies onto shore (Fig. 7; Fig. 8). The importance of wind as a driver of strandings aligns with results from previous research (Graham et al. 2001; Ferrer and Pastor 2017; Headlam et al. 2020; Macías et al. 2021; Torres-Conde et al. 2021; Bourg et al. 2022; Bourg 2024; Colaço Martins et al. 2024), and our findings highlighting currents as key drivers of offshore transport are also consistent with earlier studies (Martins 2022; Bourg 2024). We expect that this dynamic is especially robust in regions dominated by strong western boundary currents like the Gulf Stream, where the contribution of wind-forced advection, estimated at only 1.7% of wind velocity (Bourg et al. 2022), is unlikely to overcome the high velocity of the current, which advects *P. physalis* at 100% of its velocity according to the model (Eq 1).

In areas outside a strong current system like the Gulf Stream, we expect that this relationship will weaken. In the Slope Sea, we find that wind direction plays an important role in *P. physalis* transport dynamics (Fig. 9), which is likely due to a combination of direct wind-forcing and wind-induced surface currents. As a *P. physalis* colony approaches shore, surface currents decelerate and wave heights increase due to increased bottom friction and turbulence in shallower waters (Kämpf 2017; Wu et al. 2022; Zhang and Chen 2023). As a result, the ratio of wind velocity to induced surface current velocity increases, while Stokes drift becomes more influential on *P. physalis* advection. We theorize that this dynamic explains why windage and associated wave-driven processes have such a strong effect on nearshore advection and strandings (Fig. 7; Fig. 8). Further disentangling the specific contributions of windage and Stokes drift would require additional experimentation. However, given that Stokes drift is typically estimated at around 1% of wind speed (Rascle et al. 2006), we expect windage to have a relatively greater influence on *P. physalis* transport in nearshore environments than in offshore waters.

Studies of *Physalia* spp. biogeography often emphasize strandings as the endpoint of dispersal, likely due to their interactions with humans near-shore, where they can be of public health relevance (Alam and Qasim 1991; Edwards and Hessinger 2000; Cazorla-Perfetti et al. 2012; Labadie et al. 2012; Cavalcante et al. 2020). However, this emphasis on strandings complicates efforts to decouple determinants of large-scale distributions from those driving near-shore dynamics and strandings specifically. Our results suggest that onshore winds will only reliably predict strandings when *P. physalis* colonies are already present in nearshore waters (Fig. 7; Fig. 8). While wind is a strong predictor of *P. physalis* strandings at local scales and may play an important role in dispersal in regions without persistent currents, our results suggest that in the northwest Atlantic, currents are the primary driver of long-distance transport. In light of recent evidence suggesting extensive, highly interconnected *Physalia* spp. population ranges across global oceans (Church et al. 2025), we maintain that a more complete understanding of their biogeography requires targeted investigation of their open-ocean distribution and the current dynamics that drive long-distance transport in waters far from shore.

One hypothesis we considered, given the progression of strandings up the US East Coast observed in the iNaturalist data, is that the timing of regional peaks in stranding reflects delayed arrivals of *P. physalis* colonies originating in the identified surfacing region. In this scenario, colonies surfacing at the start of the year gradually disperse northward via the Gulf Stream, resulting in sequential peaks in strandings up the coastline. However, we observed rapid transport via the Gulf Stream, such that particles had populated the current past Cape Hatteras as early as January (Fig. 6A). Instead, we use the framework described above, in which ocean currents broadly distribute *P. physalis* colonies and wind facilitates coastal strandings, to explain US East Coast strandings in the three coastal regions as driven by regional wind patterns and the proximity and conditions of the Gulf Stream relative to the coastline.

We observed a substantial number of particles in our simulation that did not beach and instead dispersed eastward across the North Atlantic via the offshore region of the Gulf Stream (Fig. 6A). Although these particles were eventually removed from the simulation upon reaching the eastern boundary of our model domain, we theorize that many particles would continue dispersing east. This pattern likely reflects the underlying dynamic responsible for the genetic linkage found between some *P. physalis* specimens in the Azores and the northwest Atlantic population (Church et al. 2025).

On the east coast of Florida, the Gulf Stream flows close to shore and persistent easterly winds dominate (Peng et al. 1999), resulting in frequent strandings, proportional to the number of *P. physalis* colonies in nearby waters (Fig. 5C; Fig. 6B). From Georgia to Cape Hatteras, given the southeast-facing orientation of the coastline, we expect southerly and easterly wind patterns to drive strandings. Wind patterns in this region are variable, but strong easterly winds are relatively rare year-round (Blanton et al. 2003). Our particle-tracking simulations show that *P. physalis* often disperse into the slow moving shelf waters of the South Atlantic Bight, where they are usually retained for weeks to months (Fig. 6C). As a result, we observed a persistent nearshore population, capable of causing strandings during intermittent southerly wind events throughout the first half of the year. This explains the more diffuse temporal stranding pattern seen in the iNaturalist data from the Georgia to Cape Hatteras region compared to the other coastal regions (Fig. 2). The peak in strandings observed in June (Fig. 2B) coincides with a period when wind patterns in the region are southerly (Blanton et al. 2003). As observed in Bourg et al., 2024, strong fronts associated with western boundary currents can trap *Physalia* spp. in nearshore waters, which results in the accumulation of particles observed in the South Atlantic Bight. As the production of *P. physalis* in the winter ends and fewer colonies enter the region from the south, strandings decline through the remainder of the summer, despite continued southerly wind conditions (Fig. 2B; Fig. 5C).

North of Cape Hatteras stranding dynamics were more complex than strandings south of Cape Hatteras due to the greater distance between the Gulf Stream’s offshore path and the coastline. All simulated *P. physalis* that stranded here first dispersed northward from the offshore region of the Gulf Stream into the Slope Sea and Mid-Atlantic Bight (Fig. 6D), and we found a significant correlation between this northward exit from the Gulf Stream and prevailing southerly wind patterns (Fig. 9). Although we observed strandings in May and early June north of Cape Hatteras, when average wind patterns were predominantly northeasterly (Fig. 9), we attributed these events to particles dispersing southwestward towards the coasts of North Carolina and Virginia (Fig. 5D). We observed a similar pattern with fall strandings in Maryland, Delaware, and New Jersey (Fig. 5D). By contrast, on Long Island and in southern New England, strandings only occurred following sustained periods of southerly winds, which is due to the extensive stretch of slower-moving slope and shelf waters particles must traverse to reach the shoreline (Fig. 5D; Fig. 9).

However, this moderate correlation between southerly wind patterns and the exit of particles from the Gulf Stream does not fully explain the temporal pattern of particles exiting the Gulf Stream North of Cape Hatteras. Despite intermittent periods of strong southerly winds throughout the winter, we observed almost no movement of particles out of the Gulf Stream to the north during this season (Fig. 9). We propose that seasonal changes in the dynamics of the offshore region of the Gulf Stream may help facilitate this summertime particle exit. Our findings highlight the role of warm-core eddies in moving particles out of the Gulf Stream in the offshore region (Fig. 10). The eddy-kinetic energy of the current in this region and the formation of warm-core eddies in particular peaks during the summer (Kang et al. 2016; Silver et al. 2021). In concert, the position of the Gulf Stream’s north wall shifts north, the mean kinetic energy of the current decreases, and the frontal temperature gradient with waters to the north reduces (Fu et al. 1987; Minobe et al. 2010; Kang et al. 2016; Sánchez-Román et al. 2024). All of these conditions are known to increase the exchange of waters across current fronts and Gulf Stream surface intrusion into the Slope Sea and Mid-Atlantic Bight (Schollaert et al. 2004; Dufour et al. 2015). We theorize that this combination of seasonal changes contributes to the pronounced summer peak seen in simulated *P. physalis* exiting to the north, and subsequently strandings in New England and surrounding areas (Fig. 9). This finding aligns with the characterization of the East Australian Current in Bourg et al., 2024 as both a barrier to and facilitator of *Physalia* spp. transport towards the coast depending on the conditions of the current (Bourg 2024).

The theorized influence of seasonal dynamics of the Gulf Stream is a compelling avenue for future research, especially given the broader ecological implications. We find that the same conditions that lead to the transport of *P. physalis* out of the Gulf Stream, such as onshore surface winds and eddy formation, also broadly contribute to the intrusion of Gulf Stream water into the Slope Sea, the Mid-Atlantic Bight, and the South Atlantic Bight (Schollaert et al. 2004; Castelao 2011). Other planktonic organisms, including larvae of commercially important fish species, utilize the meanders and eddies of the Gulf Stream for transport, while frontal and ring eddies further influence regional ecology due to their distinct biological and chemical properties (Yoder et al. 1981; Pelegrí and Csanady 1991; Hare and Cowen 1996; Gaube and McGillicuddy 2017; Rypina et al. 2019). As anthropogenic climate change continues to alter Gulf Stream dynamics, we believe that further research into the relationship between *P. physalis* biogeography and these changing conditions could position *P. physalis* as a valuable indicator species, offering insights into the physical oceanography of this region and its broader ecological impacts (Gangopadhyay et al. 2019; Gai et al. 2023; Piecuch and Beal 2023; Perez et al. 2024).

## Data Availability

The code and data used to run analyses included here are available at the GitHub repository https://github.com/dunnlab/physalia-dispersal-data, commit 91f8cf9.

## Author Contributions

**RBA**: Conceptualization, methodology, software, validation, formal analysis, investigation, data curation, writing - original draft, review, and editing, visualization. **MBD**: Conceptualization, methodology, resources, writing - original draft, review, and editing, visualization, supervision, project administration. **CWD**: Conceptualization, methodology, resources, writing - original draft, review, and editing, visualization, supervision, project administration, funding acquisition. **SHC**: Conceptualization, methodology, investigation, resources, writing - original draft, review, and editing, visualization, supervision, project administration.

## Acknowledgements

We sincerely thank Dr. Amandine Schaeffer for reviewing an earlier draft of this work. We also thank Miriam Olivares for her technical support with GIS resources used in this project.

Generative AI (ChatGPT, OpenAI) was used to assist with code generation and annotation, and in condensing parts of the text.

Funding for this research was provided by the Bingham Oceanographic Collection Endowment of the Yale Peabody Museum.

## References

Alam, J. M., and R. Qasim. 1991. Toxicology of Physalia’s (Portuguese man-o’ war) venom. Pakistan Journal of Pharmaceutical Sciences 4: 159–168.

Andrews, D. G., and M. E. Mcintyre. 1978. An exact theory of nonlinear waves on a Lagrangian-mean flow. Journal of Fluid Mechanics 89: 609–646. doi:10.1017/S0022112078002773

Blanton, B. O., A. Aretxabaleta, F. E. Werner, and H. E. Seim. 2003. Monthly climatology of the continental shelf waters of the South Atlantic Bight. Journal of Geophysical Research: Oceans 108. doi:10.1029/2002JC001609

Bourg, N. 2024. Interactions between boundary currents, fronts and eddies in the Northern Current and the East Australian Current.: Transport dynamics and application to the journey of Physalia spp. PhD thesis. Université de Toulon.

Bourg, N., A. Schaeffer, P. Cetina-Heredia, J. C. Lawes, and D. Lee. 2022. Driving the blue fleet: Temporal variability and drivers behind bluebottle (Physalia physalis) beachings off Sydney, Australia D. Hyrenbach [ed.]. PLOS ONE 17: e0265593. doi:10.1371/journal.pone.0265593

Bourg, N., A. Schaeffer, A. Molcard, C. Luneau, D. E. Hewitt, and R. Chemin. 2024. Ocean wanderers: A lab-based investigation into the effect of wind and morphology on the drift of *Physalia* spp. Marine Pollution Bulletin 207: 116856. doi:10.1016/j.marpolbul.2024.116856

Burdzy, K., and Z.-Q. Chen. 2008. Discrete approximations to reflected Brownian motion. The Annals of Probability 36: 698–727. doi:10.1214/009117907000000240

Canepa, A., J. E. Purcell, P. Córdova, M. Fernández, and S. Palma. 2020. Massive strandings of pleustonic Portuguese Man-of-War (Physalia physalis) related to ENSO events along the southeastern Pacific Ocean. Latin american journal of aquatic research 48: 806–817. doi:10.3856/vol48-issue5-fulltext-2530

Castelao, R. 2011. Intrusions of Gulf Stream waters onto the South Atlantic Bight shelf. Journal of Geophysical Research: Oceans 116. doi:10.1029/2011JC007178

Cavalcante, M. M. e. S., Z. M. R. Rodrigues, R. A. Hauser-Davis, S. Siciliano, V. Haddad Júnior, and J. L. S. Nunes. 2020. Health-risk assessment of Portuguese man-of-war (*Physalia Physalis*) envenomations on urban beaches in São Luís city, in the state of Maranhão, Brazil. Revista da Sociedade Brasileira de Medicina Tropical 53: e20200216. doi:10.1590/0037-8682-0216-2020

Cazorla-Perfetti, D. J., J. Loyo, L. Lugo, M. E. Acosta, P. Morales, V. Haddad, and A. J. Rodriguez-Morales. 2012. Epidemiology of the Cnidarian Physalia physalis stings attended at a health care center in beaches of Adicora, Venezuela. Travel Medicine and Infectious Disease 10: 263–266. doi:10.1016/j.tmaid.2012.09.007

Church, S. H. and others. 2025. Population genomics of a sailing siphonophore reveals genetic structure in the open ocean. Current Biology. doi:10.1016/j.cub.2025.05.066

Clarke, A. J., and S. Van Gorder. 2018. The Relationship of Near-Surface Flow, Stokes Drift and the Wind Stress. Journal of Geophysical Research: Oceans 123: 4680–4692. doi:10.1029/2018JC014102

Colaço Martins, L., J. N. Gomes-Pereira, G. Dionísio, and J. Assis. 2024. Unravelling environmental drivers and patterns of Portuguese man o’ war (*Physalia Physalis*) blooms in two ocean regions: North Atlantic and the Southeast Pacific. Marine Pollution Bulletin 209: 117278. doi:10.1016/j.marpolbul.2024.117278

Damian-Serrano, A., S. H. D. Haddock, and C. W. Dunn. 2021. The evolution of siphonophore tentilla for specialized prey capture in the open ocean. Proceedings of the National Academy of Sciences 118: e2005063118. doi:10.1073/pnas.2005063118

Delepoulle, A., Evanmason, Clément, CoriPegliasco, A. Capet, C. Troupin, and N. Koldunov. 2022. AntSimi/py-eddy-tracker: V3.6.1. doi:10.5281/ZENODO.7197432

Dufour, C. O. and others. 2015. Role of Mesoscale Eddies in Cross-Frontal Transport of Heat and Biogeochemical Tracers in the Southern Ocean.doi:10.1175/JPO-D-14-0240.1

Edwards, L., and D. A. Hessinger. 2000. Portuguese Man-of-war (Physalia physalis) venom induces calcium influx into cells by permeabilizing plasma membranes. Toxicon 38: 1015–1028. doi:10.1016/S0041-0101(99)00213-5

European Union-Copernicus Marine Service. 2016. Global Ocean 1/12^∘^ Physics Analysis and Forecast updated Daily. doi:10.48670/MOI-00016

European Union-Copernicus Marine Service. 2021. GLOBAL OCEAN GRIDDED L4 SEA SURFACE HEIGHTS AND DERIVED VARIABLES REPROCESSED (1993- ONGOING). doi:10.48670/MOI-00148

European Union-Copernicus Marine Service. 2022. Global Ocean Hourly Sea Surface Wind and Stress from Scatterometer and Model. doi:10.48670/MOI-00305

Ferrer, L., and M. González. 2021. Relationship between dimorphism and drift in the Portuguese man-of-war. Continental Shelf Research 212: 104269. doi:10.1016/j.csr.2020.104269

Ferrer, L., and A. Pastor. 2017. The Portuguese man-of-war: Gone with the wind. Regional Studies in Marine Science 14: 53–62. doi:10.1016/j.rsma.2017.05.004

Fierro, P., L. Arriagada, A. Piñones, and J. F. Araya. 2021. New insights into the abundance and seasonal distribution of the Portuguese man-of-war *Physalia Physalis* (Cnidaria: Siphonophorae) in the southeastern Pacific. Regional Studies in Marine Science 41: 101557. doi:10.1016/j.rsma.2020.101557

Fu, L., J. Vazquez, and M. E. Parke. 1987. Seasonal variability of the Gulf Stream from satellite altimetry. Journal of Geophysical Research: Oceans 92: 749–754. doi:10.1029/JC092iC01p00749

Gai, C. and others. 2023. Heterogenous westerly shifts linked to Atlantic meridional overturning circulation slowdowns. Communications Earth & Environment 4: 325. doi:10.1038/s43247-023-00987-z

Gangopadhyay, A., G. Gawarkiewicz, E. N. S. Silva, M. Monim, and J. Clark. 2019. An Observed Regime Shift in the Formation of Warm Core Rings from the Gulf Stream. Scientific Reports 9: 12319. doi:10.1038/s41598-019-48661-9

Gaube, P., and D. J. McGillicuddy. 2017. The influence of Gulf Stream eddies and meanders on near-surface chlorophyll. Deep Sea Research Part I: Oceanographic Research Papers 122: 1–16. doi:10.1016/j.dsr.2017.02.006

Graham, W. M., F. Pagès, and W. M. Hamner. 2001. A physical context for gelatinous zooplankton aggregations: A review. Hydrobiologia 451: 199–212. doi:10.1023/A:1011876004427

Groeskamp, S., J. H. LaCasce, T. J. McDougall, and M. Rogé. 2020. Full-Depth Global Estimates of Ocean Mesoscale Eddy Mixing From Observations and Theory. Geophysical Research Letters 47: e2020GL089425. doi:10.1029/2020GL089425

Hare, J. A., and R. K. Cowen. 1996. Transport mechanisms of larval and pelagic juvenile bluefish (*Pomatomus Saltatrix*) from South Atlantic Bight spawning grounds to Middle Atlantic Bight nursery habitats. Limnology and Oceanography 41: 1264–1280. doi:10.4319/lo.1996.41.6.1264

Headlam, J., K. Lyons, J. Kenny, E. S. Lenihan, D. Quigley, W. Helps, M. M. Dugon, and T. K. Doyle. 2020. Insights on the origin and drift trajectories of Portuguese man of war (Physalia physalis) over the Celtic Sea shelf area. Estuarine, Coastal and Shelf Science 246. doi:10.1016/j.ecss.2020.107033

Helm, R. R. 2021. The mysterious ecosystem at the ocean’s surface. PLOS Biology 19: e3001046. doi:10.1371/journal.pbio.3001046

Hilton, A. 2025. South Florida’s East coast and its unruly inhabitants: Portuguese men-of-war. WLRN.

Hogg, N. G. 1992. On the transport of the gulf stream between cape hatteras and the grand banks. Deep Sea Research Part A. Oceanographic Research Papers 39: 1231–1246. doi:10.1016/0198-0149(92)90066-3

Hyun, S. and others. 2022. Ocean mover’s distance: Using optimal transport for analysing oceanographic data. Proceedings of the Royal Society A: Mathematical, Physical and Engineering Sciences 478: 20210875. doi:10.1098/rspa.2021.0875 iNaturalist. iNaturalist.

iNaturalist Community. 2024. Observations of Physalia physalis from the northwest Atlantic observed from 2000–01-07 to 2024-10-06.

Iosilevskii, G., and D. Weihs. 2009. Hydrodynamics of sailing of the Portuguese man-of-war *Physalia Physalis*. Journal of The Royal Society Interface 6: 613–626. doi:10.1098/rsif.2008.0457

Kämpf, J. 2017. Wind-Driven Overturning, Mixing and Upwelling in Shallow Water: A Nonhydrostatic Modeling Study. Journal of Marine Science and Engineering 5: 47. doi:10.3390/jmse5040047

Kang, D., E. N. Curchitser, and A. Rosati. 2016. Seasonal Variability of the Gulf Stream Kinetic Energy. Journal of Physical Oceanography 46: 1189–1207. doi:10.1175/JPO-D-15-0235.1

Kehl, C., P. D. Nooteboom, M. L. A. Kaandorp, and E. Van Sebille. 2023. Efficiently simulating Lagrangian particles in large-scale ocean flows — Data structures and their impact on geophysical applications. Computers & Geosciences 175: 105322. doi:10.1016/j.cageo.2023.105322

Kitanidis, P. K. 1994. Particle-tracking equations for the solution of the advection-dispersion equation with variable coefficients. Water Resources Research 30: 3225–3227. doi:10.1029/94WR01880

Klein, A. 2023. Swimmers beware: Portuguese man-of-wars washing up on New England beaches. NBC Boston.

Labadie, M. and others. 2012. Portuguese man-of-war (Physalia physalis) envenomation on the Aquitaine Coast of France: An emerging health risk. Clinical Toxicology (Philadelphia, Pa.) 50: 567–570. doi:10.3109/15563650.2012.707657

Lee, D., A. Schaeffer, and S. Groeskamp. 2021. Drifting dynamics of the bluebottle (*Physalia Physalis*). Ocean Science 17: 1341–1351. doi:10.5194/os-17-1341-2021

Macías, D., L. Prieto, and E. García-Gorriz. 2021. A model-based management tool to predict the spread of Physalia physalis in the Mediterranean Sea. Minimizing risks for coastal activities. Ocean & Coastal Management 212: 105810. doi:10.1016/j.ocecoaman.2021.105810

Martins, L. C. 2022. Modelling the occurrence of Physalia physalis in the North Atlantic Ocean at different spatial and temporal scales. PhD thesis. Universidade Do Algarve.

Mason, E., A. Pascual, and J. C. McWilliams. 2014. A New Sea Surface Height–Based Code for Oceanic Mesoscale Eddy Tracking. Journal of Atmospheric and Oceanic Technology 31: 1181–1188. doi:10.1175/JTECH-D-14-00019.1

Meinen, C. S., and D. S. Luther. 2016. Structure, transport, and vertical coherence of the Gulf Stream from the Straits of Florida to the Southeast Newfoundland Ridge. Deep Sea Research Part I: Oceanographic Research Papers 112: 137–154. doi:10.1016/j.dsr.2016.03.002

Minobe, S., M. Miyashita, A. Kuwano-Yoshida, H. Tokinaga, and S.-P. Xie. 2010. Atmospheric Response to the Gulf Stream: Seasonal Variations. doi:10.1175/2010JCLI3359.1

Mohtar, S. E., I. Hoteit, O. Knio, L. Issa, and I. Lakkis. 2018. Lagrangian tracking in stochastic fields with application to an ensemble of velocity fields in the Red Sea. Ocean Modelling 131: 1–14. doi:10.1016/j.ocemod.2018.08.008

Muller-Karger, F. E. and others. 2015. Natural variability of surface oceanographic conditions in the offshore Gulf of Mexico. Progress in Oceanography 134: 54–76. doi:10.1016/j.pocean.2014.12.007

Munro, C., Z. Vue, R. R. Behringer, and C. W. Dunn. 2019. Morphology and development of the Portuguese man of war, Physalia physalis. Scientific Reports 9: 15522. doi:10.1038/s41598-019-51842-1

Naisbett-Jones, L. C., N. F. Putman, J. F. Stephenson, S. Ladak, and K. A. Young. 2017. A Magnetic Map Leads Juvenile European Eels to the Gulf Stream. Current Biology 27: 1236–1240. doi:10.1016/j.cub.2017.03.015

Natural Earth contributors. Made with Natural Earth. Free vector and raster map data @ naturalearthdata.com.

Oguchi, K., G. Yamamoto, H. Kohtsuka, and C. W. Dunn. 2024. Physalia gonodendra are not yet sexually mature when released. Scientific Reports 14: 23011. doi:10.1038/s41598-024-73611-5

Pegliasco, C., A. Delepoulle, E. Mason, R. Morrow, Y. Faugère, and G. Dibarboure. 2022. META3.1exp: A new global mesoscale eddy trajectory atlas derived from altimetry. Earth System Science Data 14: 1087–1107. doi:10.5194/essd-14-1087-2022

Pelegrí, J. L., and G. T. Csanady. 1991. Nutrient transport and mixing in the Gulf Stream. Journal of Geophysical Research: Oceans 96: 2577–2583. doi:10.1029/90JC02535

Peng, G., C. N. K. Mooers, and H. C. Graber. 1999. Coastal Winds in South Florida.

Perez, E., M. Andres, and G. Gawarkiewicz. 2024. Is the Regime Shift in Gulf Stream Warm Core Rings Detected by Satellite Altimetry? An Inter-Comparison of Eddy Identification and Tracking Products. Journal of Geophysical Research: Oceans 129: e2023JC020761. doi:10.1029/2023JC020761

Piecuch, C. G., and L. M. Beal. 2023. Robust Weakening of the Gulf Stream During the Past Four Decades Observed in the Florida Straits. Geophysical Research Letters 50: e2023GL105170. doi:10.1029/2023GL105170

Prieto, L., D. Macías, A. Peliz, and J. Ruiz. 2015. Portuguese Man-of-War (Physalia physalis) in the Mediterranean: A permanent invasion or a casual appearance? Scientific Reports 5: 11545. doi:10.1038/srep11545

Pugh, P. R. 2019. A history of the sub-order Cystonectae (Hydrozoa: Siphonophorae). Zootaxa 4669. doi:10.11646/zootaxa.4669.1.1

QGIS Development Team. 2009. QGIS Geographic Information System.

Rascle, N., F. Ardhuin, and E. A. Terray. 2006. Drift and mixing under the ocean surface: A coherent one-dimensional description with application to unstratified conditions. Journal of Geophysical Research: Oceans 111. doi:10.1029/2005JC003004

Ross, O. N., and J. Sharples. 2004. Recipe for 1-D Lagrangian particle tracking models in space-varying diffusivity. Limnology and Oceanography: Methods 2: 289–302. doi:10.4319/lom.2004.2.289

Rypina, I. I., K. Chen, C. M. Hernández, L. J. Pratt, and J. K. Llopiz. 2019. Investigating the suitability of the Slope Sea for Atlantic bluefin tuna spawning using a high-resolution ocean circulation model. ICES Journal of Marine Science 76: 1666–1677. doi:10.1093/icesjms/fsz079

Sánchez-Román, A., F. Gues, R. Bourdalle-Badie, M.-I. Pujol, A. Pascual, and M. Drévillon. 2024. Changes in the Gulf Stream path over the last 3 decades. State of the Planet 4-osr8: 1–11. doi:10.5194/sp-4-osr8-4-2024

Schollaert, S. E., T. Rossby, and J. A. Yoder. 2004. Gulf Stream cross-frontal exchange: Possible mechanisms to explain interannual variations in phytoplankton chlorophyll in the Slope Sea during the SeaWiFS years. Deep Sea Research Part II: Topical Studies in Oceanography 51: 173–188. doi:10.1016/j.dsr2.2003.07.017

Silver, A., A. Gangopadhyay, G. Gawarkiewicz, E. N. S. Silva, and J. Clark. 2021. Interannual and seasonal asymmetries in Gulf Stream Ring Formations from 1980 to 2019. Scientific Reports 11: 2207. doi:10.1038/s41598-021-81827-y

Stommel, H. 1948. The westward intensification of wind-driven ocean currents. Eos, Transactions American Geophysical Union 29: 202–206. doi:10.1029/TR029i002p00202

Storer, B. A., M. Buzzicotti, H. Khatri, S. M. Griffies, and H. Aluie. 2023. Global cascade of kinetic energy in the ocean and the atmospheric imprint. Science Advances 9: eadi7420. doi:10.1126/sciadv.adi7420

Taylor, J. C., J. M. Miller, L. J. Pietrafesa, D. A. Dickey, and S. W. Ross. 2010. Winter winds and river discharge determine juvenile southern flounder (*Paralichthys Lethostigma*) recruitment and distribution in North Carolina estuaries. Journal of Sea Research 64: 15–25. doi:10.1016/j.seares.2009.09.006

Tewari, A., T. Yin, G. Cazenavette, S. Rezchikov, J. B. Tenenbaum, F. Durand, W. T. Freeman, and V. Sitzmann. 2023. Diffusion with Forward Models: Solving Stochastic Inverse Problems Without Direct Supervision. doi:10.48550/ARXIV.2306.11719

Torres Conde, E. G., and R. E. Rodríguez Martínez. 2023. The Largest Historical Stranding of Physalia physalis Recorded. doi:10.20944/preprints202307.1676.v1

Torres-Conde, E. G. 2022. Is simultaneous arrival of pelagic Sargassum and Physalia physalis a new threat to the Atlantic coasts? Estuarine, Coastal and Shelf Science 275: 107971. doi:10.1016/j.ecss.2022.107971

Torres-Conde, E. G., B. Martínez-Daranas, and L. Prieto. 2021. The Havana littoral, an area of distribution for Physalia physalis in the Atlantic Ocean. Regional Studies in Marine Science 44: 101752. doi:10.1016/j.rsma.2021.101752

Totton, A. K. 1960. Studies on Physalia physalis (L.). Pt. 1 Natural history and morphology. Discovery Reports 30: 301–368.

Totton, A. K., and G. O. Mackie. 1956. Dimorphism in the Portuguese Man-of-War. Nature 177: 290–290. doi:10.1038/177290b0

Wittenberg, J. B. 1960. The Source of Carbon Monoxide in the Float of the Portuguese Man-of-War, *Physalia Physalis* L. Journal of Experimental Biology 37: 698–705. doi:10.1242/jeb.37.4.698

Wu, T., B. Qin, A. Huang, Y. Sheng, S. Feng, and C. Casenave. 2022. Reconsideration of wind stress, wind waves, and turbulence in simulating wind-driven currents of shallow lakes in the Wave and Current Coupled Model (WCCM) version 1.0. Geoscientific Model Development 15: 745–769. doi:10.5194/gmd-15-745-2022

Yang, B., F. Gomez, C. Schmid, and M. Baringer. 2022. In situ estimates of net primary production in the open-ocean Gulf of Mexico. Limnology and Oceanography Letters 7: 427–434. doi:10.1002/lol2.10270

Yoder, J. A., L. P. Atkinson, T. N. Lee, H. H. Kim, and C. R. McClain. 1981. Role of Gulf Stream frontal eddies in forming phytoplankton patches on the outer southeastern shelf. Limnology and Oceanography 26: 1103–1110. doi:10.4319/lo.1981.26.6.1103

Zhang, C., and L. Chen. 2023. A review of wind-driven hydrodynamics in large shallow lakes: Importance, process-based modeling and perspectives. Cambridge Prisms: Water 1: e16. doi:10.1017/wat.2023.14

