## Supporting Information for "From surfacing to stranding: The origins and dispersal dynamics of a neustonic siphonophore"

River B. Abedon[1,*], Mary Beth Decker[1], Casey W. Dunn[1], Samuel H. Church[1,2,*]

1. Yale University, Department of Ecology and Evolutionary Biology, New Haven, Connecticut, USA
2. New York University, Department of Biology, New York, New York, USA

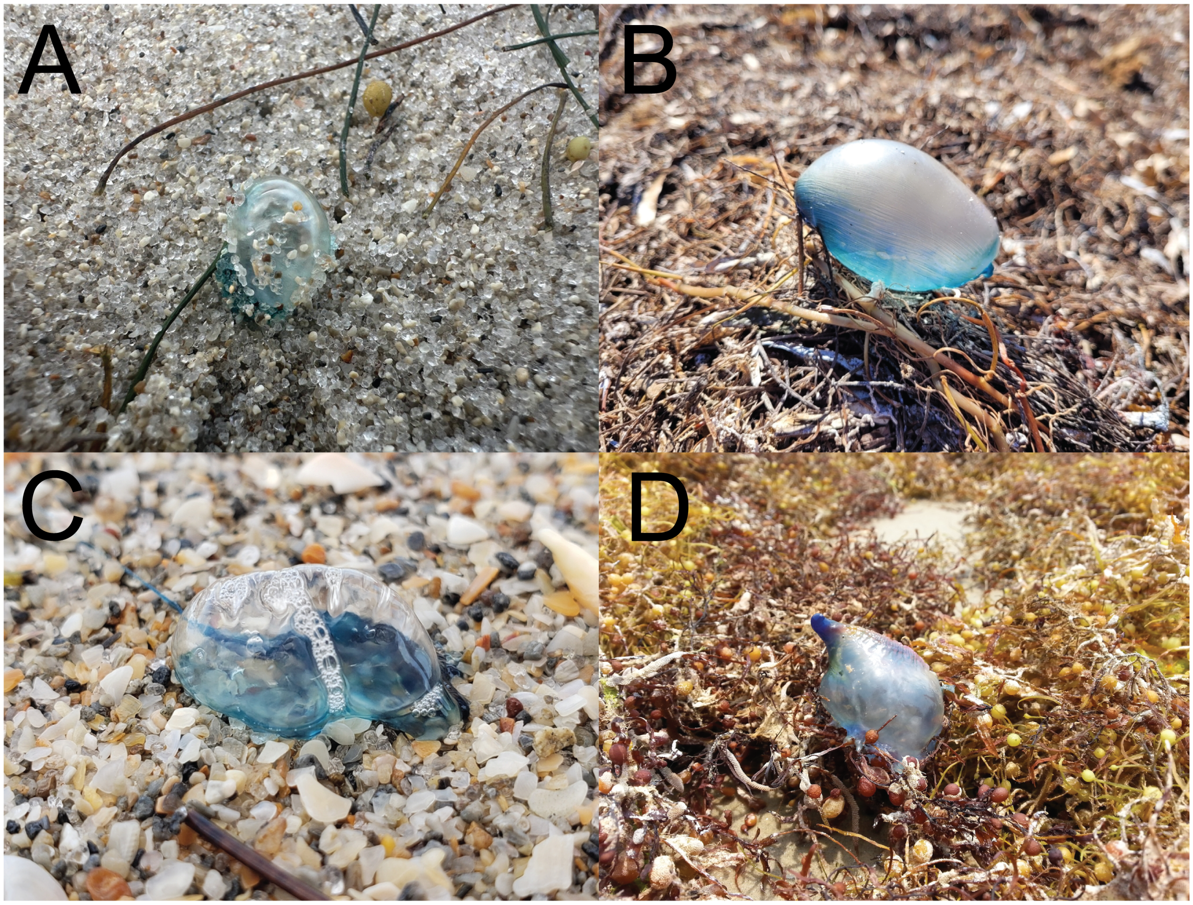


**Figure S1**: **Examples of iNaturalist images classified as juveniles.** Images from iNaturalist *P. physalis* observations demonstrating diagnostic characteristics used to classify specimens as juvenile. The colonies shown have a small float, highly reduced or absent crest/sail, and few visible zooids. The specimens also have clear, unobscured morphology and appear undamaged. Observations by Nick Ellis, Dylan Friedman, Oliver G Jones, wanderingeden licensed under CC BY-NC 4.0.

The primary diagnostic used to classify imaged *Physalia physalis* specimens as juveniles was the presence of few colony bodies (zooids). As *Physalia* colonies mature, new zooids are added and existing zooids grow in size (Munro et al. 2019). Therefore, a juvenile colony is characterized by a limited number of small zooids on the ventral and posterior side of the float (S1A; S1D). Juvenile *P. physalis* colonies are also expected to have smaller floats than mature colonies (S1), as juveniles surface once their float inflates sufficiently to achieve positive buoyancy and will continue to grow and inflate as the colony matures (Munro et al. 2019). Float size was inferred from iNaturalist images using approximate reference objects such as seaweed (S1B; S1C), coarse sand (S1A; S1C), shells, or parts of the observer's body, where visible. In addition to these characters, specimens were considered juvenile when they had no visible tentacles or only one relatively small tentacle (S1), and we considered any specimen with several tentacles or particularly large tentacles mature, as well as any specimens with prominent and well-developed crests. All observations classified as juveniles were re-evaluated to ensure they align with the diagnostics described above, resulting in the removal of two observations from the final dataset that were found to be misclassifications on the first pass.

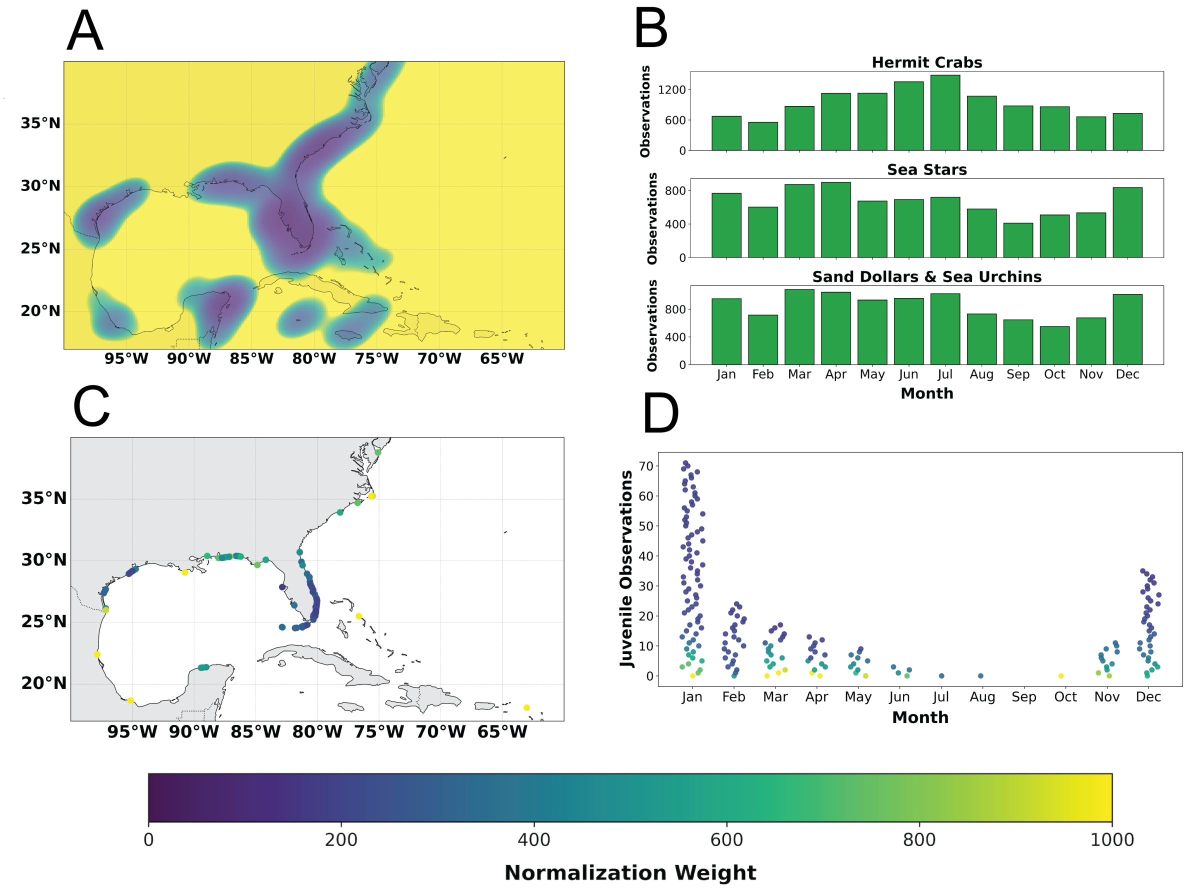


**Figure S2**: **Normalizing juvenile distribution for iNaturalist user effort.** (A) Heatmap of geographic dimension of normalization weights calculated using equation S3. Normalization weights range from 0 to 1000. Yellow regions show areas categorized as "low effort/high weight" and assigned the highest normalization weight of 1000 (B) Monthly iNaturalist observations of three taxa used as proxies for user effort (C) Map of juvenile *P. physalis* observations color-coded according to normalization weight. Observations in regions and times of low user effort were assigned higher weights than those in regions and times of high user effort (D) Monthly observations of juvenile *P. physalis* color-coded according to normalization weight

To represent iNaturalist user effort, we used all observations of the taxa Paguroidea (superfamily), Asteroidea (class), and Echinoidea (class) recorded within the region of interest through 19 September 2024, the date the data were accessed, for a total of 29,782 observations in the effort dataset. We chose these groups as proxies for user effort for two main reasons. First, members of these clades are conspicuous, widespread in the region of interest, and commonly observed on beaches, which makes them likely to be recorded on iNaturalist in the same areas *P. physalis* is commonly found. Second, by using higher taxonomic ranks, the selected groups encompass broad taxonomic diversity, which reduces the likelihood that the data reflects species-specific patterns of abundance driven by seasonality or anything other than observation effort.

In order to accurately represent the temporal dimension of the iNaturalist effort data, we converted each observation's date into a numerical Day-of-Year value ranging from 1 to 366. To represent the cyclical nature of annual patterns in effort, we then transformed this value into two circular variables using the equations,

$$\text{DayOfYear\_sin} = \sin\left(\frac{2\pi \cdot \text{DayOfYear}}{365}\right) \quad \text{(S1)}$$

$$\text{DayOfYear\_cos} = \cos\left(\frac{2\pi \cdot \text{DayOfYear}}{365}\right) \quad \text{(S2)}$$

This transformation creates a circular representation of the Day-of-Year value, preserving the continuity between the end and beginning of a year.

We then used the iNaturalist effort data, comprising longitude, latitude, DayofYear_sin, and DayofYear_cos, to generate a four-dimensional kernel-density estimation (KDE) using the 'gaussian_kde' function from the SciPy statistics package. This KDE allowed us to estimate iNaturalist user effort for any given location and time throughout the year.

We applied a similar process to the juvenile observation data, converting the observation dates into Day-of-Year values. We evaluated the effort KDE at each of the 194 juvenile observation locations and times, and computed a normalization weight using the equation,

$$\text{Normalization Weight} = \min\left(\frac{1}{\text{Effort Value}},\ 1000\right) \quad \text{(S3)}$$

where the normalization weight for each juvenile observation is equal to the inverse of the effort value, with a maximum limit of 1000 (S2). We chose the normalization weight to be inversely proportional to the iNaturalist effort value such that juveniles observed in high-effort areas were weighted less than those in low-effort areas. The limit of 1000 addressed cases in which very low effort values resulted in excessively high normalization weights, resulting in over-normalization. We assigned these observations a normalization weight of 1000, which we chose to categorically represent "low effort/high weight" areas and times (S2). Due to geographic limitations in the effort dataset, we assigned a single outlier observation from St. Martin a normalization weight of 1000, because it was located beyond the effort dataset boundary (S2A; S2C).

We generated a four-dimensional KDE of juvenile *P. physalis* observations by applying the 'gaussian_kde' function to spatial coordinates and circular Day-of-Year values (Eq. S1-S2) and adjusting for iNaturalist effort using the assigned normalization weights. To initialize the particle tracking simulation, we sampled from this KDE, converting the sampled DayOfYear_sin and DayOfYear_cos values back into a single value using the equation,

$$\text{DayOfYear} = \frac{\arctan 2(\text{DayOfYear\_sin},\ \text{DayOfYear\_cos}) \times 365}{2\pi} \quad \text{(S4)}$$

which we stored alongside the particle's location, delaying advection until the assigned Day-of-Year was reached during the simulation.

Because juveniles were observed on shorelines, resulting in significant overlap of the juvenile KDE with land areas, we implemented two methods to ensure starting points accurately represented offshore surfacing locations suitable for the particle tracking model. First, we used a landmask raster to identify and discard points generated on land. We repeated the process until 10,000 valid points were generated in water. Second, we identified all of the points within 50 km of the shoreline and used a breadth-first search algorithm to move the particle to the nearest location outside of this 50 km buffer.

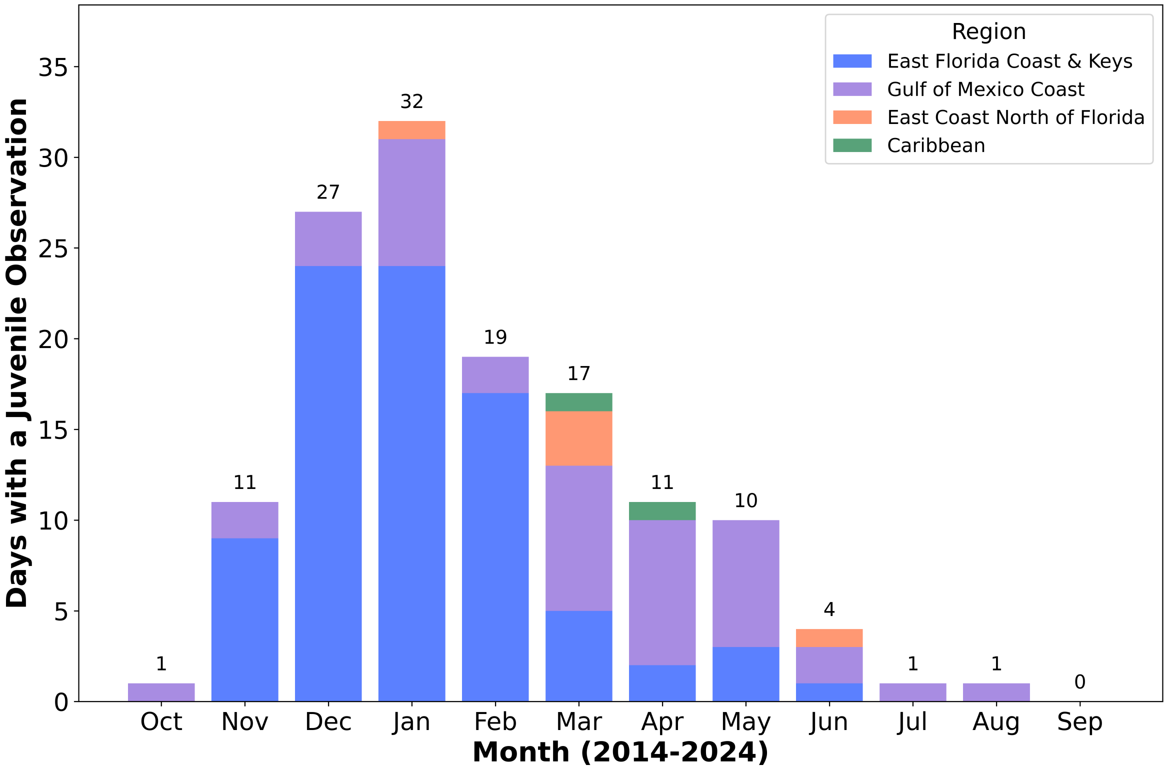


**Figure S3**: **Days with at least one juvenile *P. physalis* observation.** iNaturalist observations of juveniles are collapsed such that multiple observations of juveniles occurring on the same day in the same region are considered one data point

Munro, C., Z. Vue, R. R. Behringer, and C. W. Dunn. 2019. Morphology and development of the Portuguese man of war, Physalia physalis. Scientific Reports **9**: 15522. doi:10.1038/s41598-019-51842-1
